# Oncogene EVI1 Drives Acute Myeloid Leukemia Via a Targetable Interaction with CTBP2

**DOI:** 10.1101/2023.07.11.548358

**Authors:** Dorien Pastoors, Marije Havermans, Roger Mulet-Lazaro, Duncan Brian, Willy Noort, Julius Grasel, Remco Hoogenboezem, Leonie Smeenk, Jeroen A.A. Demmers, Michael D. Milsom, Tariq Enver, Richard WJ Groen, Eric Bindels, Ruud Delwel

## Abstract

Acute myeloid leukemia (AML) driven by the activation of EVI1 due to chromosome 3q26/MECOM rearrangements is incurable. Since transcription factors like EVI1 are notoriously hard to target, insight into the mechanism by which EVI1 drives myeloid transformation could provide alternative avenues for therapy. Applying protein folding predictions combined with proteomics technologies, we demonstrate that interaction with CTBP1 and CTBP2 via a single PLDLS motif in EVI1 is indispensable for leukemic transformation. Furthermore, we show that a 4x PLDLS repeat construct outcompetes binding of EVI1 to CTBP1 and CTBP2 and thereby inhibits proliferation of 3q26/MECOM rearranged AML both in in vitro and in xenotransplant models. This proof-of-concept study opens the possibility to therapeutically target one of the most incurable forms of AML with specific EVI1-CTBP inhibitors. This has important implications for other tumour types with aberrant expression of EVI1 as well as for cancers transformed by distinct CTBP-dependent oncogenic transcription factors.

## Introduction

Aberrant activation of EVI1 in acute myeloid leukemia (AML) is associated with poor treatment response and diminished survival (1). EVI1-expressing leukemias include those with 3q26/MECOM (MDS1 and EVI1 complex locus, from which EVI1 is transcribed) or 11q23/MLL rearrangements. In the case of 3q26/MECOM rearrangements, active enhancers translocate towards the MECOM locus (2–5). In approximately 40% of 11q23/MLL-rearranged AML patients, EVI1 is overexpressed by a mechanism that is incompletely understood (6). Irrespective of the mechanism of activation, EVI1 is essential for the survival, proliferation and the undifferentiated phenotype of those AML cells (3). Consequently, eliminating EVI1 or interfering with its function may constitute an effective therapeutic strategy for these aggressive forms of AML.

EVI1 encodes a nuclear protein containing two DNA-binding zinc-finger domains, which is an essential transcriptional regulator that has important repressive functions (7, 8). In addition to EVI1, a long isoform MDS1-EVI1 (PRDM3) is also transcribed from this locus. The key difference between these two isoforms is that MDS-EVI1, contains a SET domain implicated in histone methylation (9). However, MDS1-EVI1 is not or hardly expressed in 3q26/MECOM-rearranged AML (2). This raises the question of how EVI1 represses transcription and causes leukemic transformation. It is likely that EVI1 recruits other proteins to its binding sites in the genome to exert its repressive effect. Exact knowledge about those partners and the mechanism of interaction may be used to target EVI1-driven AML.

Several EVI1 interaction partners have been identified previously in various experimental systems, including CTBP1/2, MBD3, EHMT2, HDAC1/2, SUV39H1, and SMAD3 (10–16). In this study we investigated which proteins bind to EVI1 in AML cells and are essential for leukemic transformation. We identified CTBP1 and 2 as the most enriched EVI1-binding partners, and showed that the interaction between EVI1 and CTBP2 is essential for leukemic cell proliferation. This interaction depends on a PLDLS domain within EVI1. A PLDLS-containing repeat construct outcompetes EVI1-CTBP interaction and reverses the leukemic phenotype. This proof-of-concept-study demonstrates the possibility of targeting a subset of high-risk AML by interfering between EVI1 and its protein partners.

## Results

### EVI1 is essential for CTBP2 recruitment to chromatin in inv(3) AML

EVI1 has been reported to interact with a wide range of repressive transcriptional and epigenetic regulators, but it is unclear which of those are consistently required for EVI1-mediated transformation. To identify EVI1-binding proteins, we performed EVI1 immunoprecipitation (IP) followed by mass spectrometry (MS) in nuclear lysates (EVI1-IP/MS) from EVI1-overexpressing, 3q26/MECOM-rearranged (inv(3)) MUTZ3 cells. In this way, we identified EVI1 binding partners previously reported in multiple experiments on BioGRID, including CTBP1/2, HDAC2, RBBP4, MTA and RCOR1, of which CTBP1 and CTBP2 were most enriched compared to IgG control (10–12, 16, 22, 34, 35) (Fig. 1A). We confirmed the reciprocal interaction between the two by EVI1- or CTBP2 IP, followed by western blot (WB) for CTBP2 (EVI1-IP/CTBP2-WB) or EVI1 (CTBP2-IP/EVI1-WB), respectively (Fig S1A). We performed enrichment analysis with EnrichR on the 460 significantly co-immunoprecipitated proteins using protein complexes from the CORUM database (Fig. 1B, Fig S1B) (23). CTBP1 and CTBP2 have a central position in the network visualisation of enriched proteins connected and clustered by their co-occurrence in protein complexes (Fig. 1B, Fig S1B). To validate our findings in a different model, we performed streptavidin-based protein precipitation on murine EVI1 fused to a biotag biotinylated in vivo by E. coli biotin ligase BirA, in the murine leukemia cell line NFS78 (36) (Fig S1C). Here, we again identified CTBP1/2 as most enriched compared to IgG control (Fig S1D). Since CTBP1 and CTBP2 are highly homologous at the protein level, and both equally co-precipitated with EVI1, we decided to focus on CTBP2 to represent EV1-CTBP interaction. ChIP-seq using chromatin from MUTZ3 cells (Fig. 1C) showed a strong reciprocal correlation between EVI1 and CTBP2 binding sites (Fig. 1D, Fig. S2A, Fig. S2B). This was much higher than the correlation of either EVI1 or CTBP2 with other factors such as p300, MYB or RUNX1 (Fig. 1D, Fig. S2A). This demonstrates a strong and consistent interaction between EVI1 and CTBP2 at chromatin in AML with inv(3). Furthermore, knock-down of EVI1 (Fig. S2C) resulted in a complete loss of CTBP2 signal in ChIP-seq (Fig. 1E), demonstrating EVI1 is critical for CTBP2 to engage with chromatin in these cells.

**Figure 1:**
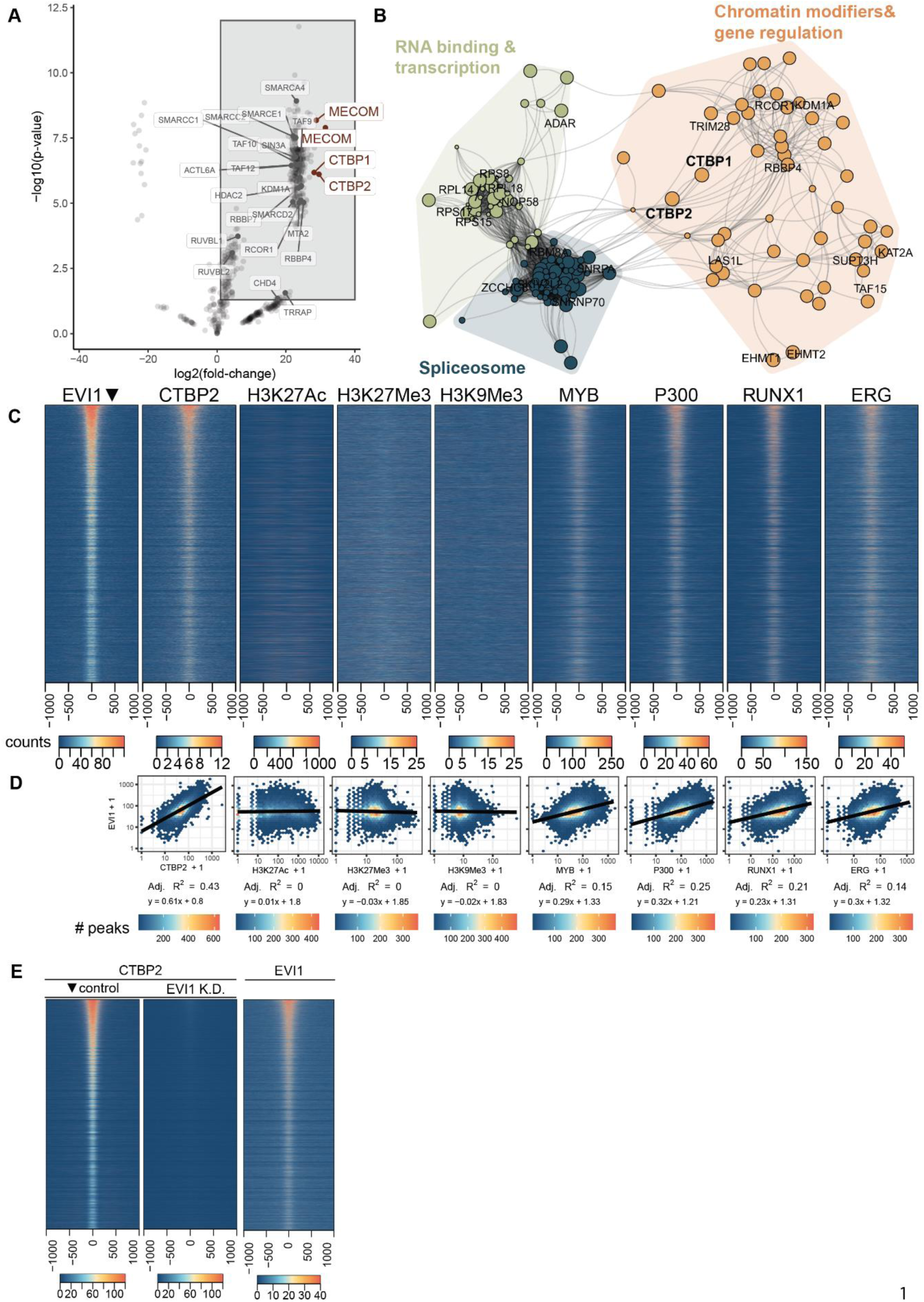
CTBP1 and CTBP2 bind EVI1. A. Differential enrichment of proteins binding to EVI1 as determined by an EVI1 specific immunoprecipitation (IP) compared to a non-specific IgG IP followed by mass spectrometry (MS) in MUTZ3 nuclear protein extracts (n = 3 for each group, significance threshold log2FC > 1 and p-value < 0.05). Top-4 enriched gene identifiers are highlighted in red (MECOM (EVI1), CTBP1 and CTBP2). In addition, several previously identified MECOM interactors are shown in grey (BioGrid). B. Network plot of protein complex enrichment analysis of the proteins co-precipitated with EVI1 IP in MUTZ3 cells. The connections (edges) between proteins (nodes) were weighted based on their co-occurrence in CORUM complexes, and the layout of the graph was determined by the Fruchterman-Reingold algorithm. Node colour represents the cluster assigned to a protein as determined by the Louvain community detection algorithm; dot size represents the fold enrichment in the IP. Font size of CTBP1 and CTBP2 was adjusted in bold manually, and coloured areas and their corresponding labels were added manually based on the common biochemical functions of the complexes in each cluster. The full plot with all labels is presented in Fig. S1A. C. Heatmaps of ChIP-Seq experiments in MUTZ3 cells using antibodies directed to the indicated transcription factors or histone modifications. Ranking was based on the ChIP-seq EVI1 signal (leftmost panel) showing signal intensity of indicated ChIP-seq tracks in a ±1000 bp region centered on EVI1 peaks. D. Quantification of the heatmap in panel Fig. 1C. Normalised EVI1 signal is plotted versus the normalised reads in the indicated ChIP-seq-tracks, for all EVI1 peaks with a window of ±1000bp. Correlation coefficients and linear regression equations are shown for log10-transformed data with a pseudo count of 1. E. Heatmap of CTBP2 ChIP-seq data following dox-inducible shRNA mediated knockdown of EVI1 (48hrs), in MOLM1 cells. On the right, EVI1 heatmap in unperturbed MOLM1 cells. Tracks are ranked on peaks in the CTBP2 control track (leftmost panel).

### A single PLDLS domain in EVI1 is critical for CTBP2 binding

To study whether the interaction between EVI1 and CTBP2 is required for transformation, we first set out to determine which amino acids in EVI1 bind CTBP2. Computational modelling of protein-protein interfaces with AlphaFold (37) predicted interaction between EVI1 and CTBP2 via a PLDLS site in EVI1 (Fig 2A, Fig 2B) with R43, H69 and K71 in CTBP2 (Fig 2B). In line with this prediction, CTBP proteins have been reported to bind to PXDLX motifs, of which there are two in EVI1, i.e. a PFDLT and a PLDLS motif (38) (Fig 2B, Fig 2C). To experimentally determine whether it is indeed the PLDLS site in EVI1 that interacts with CTBP2, distinct FLAG-tagged EVI1 mutants with deletions of PFDLT, PLDLS or both were introduced into HEK293T cells. Upon FLAG-IP, only the proteins with an intact PLDLS site were able to bind CTBP2 (Fig 2C). In addition, FLAG-tagged full length EVI1 in which PLDLS was mutated into PLASS also lost CTBP2 interaction (Fig 2C). This is in line with a reduced interaction frequency predicted by AlphaFold for a heterodimer of CTBP2 and PLASS-mutant EVI1 (Fig 2B). An EVI1 construct in which PFDLT was mutated into PFAST was still able to bind CTBP2, confirming that only PLDLS is essential for CTBP2 binding (Fig 2C).

**Figure 2:**
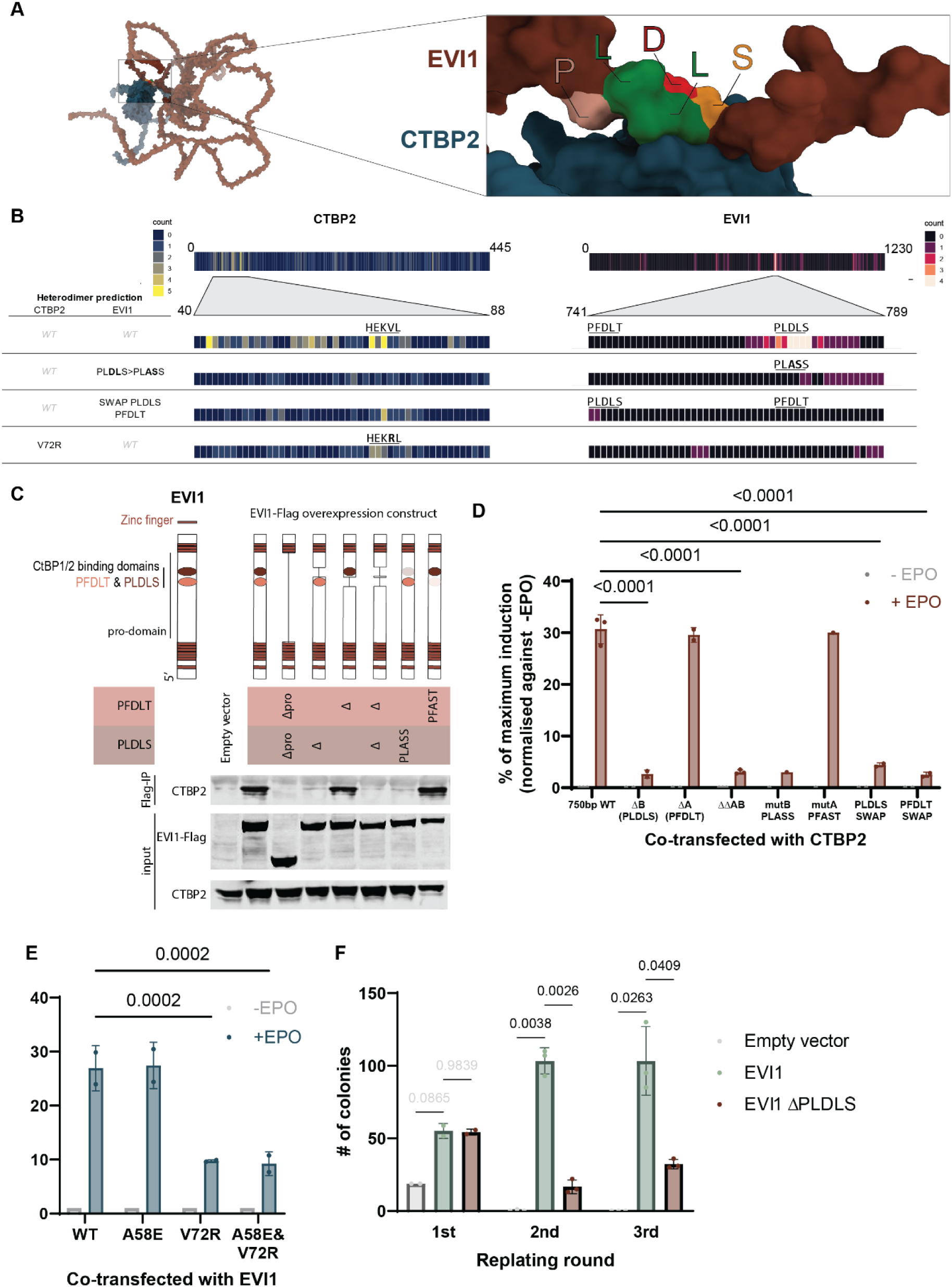
Identification of a competitive inhibitor of EVI1-CTBP2 interaction. A. EVI1-CTBP2 interaction as predicted by AlphaFold and visualized with ChimeraX. EVI1 is shown in maroon, with the PLDLS residues in different colours according to their biochemical properties. CTBP2 is depicted in dark blue. Fig 2B. Frequency of the involvement of individual residues in EVI1 and CTBP2 in the interaction between these two proteins as predicted by AlphaFold and quantified in ChimeraX (using the “interfaces” command with default parameters) for all 25 structures per heterodimer prediction. Fig 2C. Immunoprecipitation of overexpressed EVI1-FLAG with anti-FLAG antibody, followed by western blot for CTBP2 and FLAG in lysates from HEK293T cells transfected with WT or mutated EVI1-FLAG. A schematic diagram is added above the blot to indicate the location of introduced deletions/mutations. Fig 2D. MAPPIT assay in HEK293T cells to measure EVI1-CTBP interaction. Mutations introduced in EVI1 are similar to those in panel 2C. Reporter induction is normalised to –EPO (not induced) and maximum induction with RNF as positive control. Statistical significance is determined with a mixed-effects model (significance shown if P<0.05 and N > 1) in 1-3 independent experiments (all independent experiments are performed in triplicate); mean + SD is shown. Fig 2E. MAPPIT assay in HEK293T cells to measure EVI1-CTBP2 interaction, with mutations disrupting residues in CTBP2. Those residues have been previously reported to be important for target protein binding via PXDLS motifs (*49*) (n = 2 in triplicate). Reporter induction is normalised to –EPO (not induced) and maximum induction with RNF as positive control. Statistical significance is determined with a two-way ANOVA (significance shown if P<0.05) in 2 independent experiments (all independent experiments are performed in triplicate); mean + SD is shown. Fig 2F. Colony formation of mouse bone marrow cells transduced with empty vector, full-length EVI1 or EVI1 without PLDLS domain. Statistical significance was determined with a two-way ANOVA with EVI1 as reference (all comparisons shown). Experiment performed in triplicate.

To enable rapid and direct measurement of interaction between EVI1 and CTBP2, and to study the effect of different mutations, we adapted and applied the luciferase-based Mammalian protein-protein interaction trap (MAPPIT) assay (Fig. S3A) (39). In short, this assay uses JAK-STAT signalling downstream of a modified EPO receptor to quantify protein-protein interaction. EPO can activate a STAT3-responsive luciferase reporter only if there is interaction between bait and prey proteins, in this case EVI1 and CTBP2 (Fig. S3A). In the MAPPIT assay, the target protein-protein interaction takes place in the cytosol, which means that most other nuclear proteins are absent and any interaction measured is likely to occur directly between the two studied proteins. With this assay, we confirmed that only by deleting or mutating the PLDLS site the luciferase activity was abolished (Fig 2D). In addition, moving the PLDLS site to the PFDLT location and vice versa was not sufficient to rescue the interaction between EVI1 and CTBP2. This is in line with predictions by AlphaFold (Fig 2B, Fig 2D. These results indicate that the PLDLS residues themselves, as well as some surrounding amino acids, are likely essential for the interaction between EVI1 and CTBP2. Within CTBP2, EVI1 is predicted to interact mostly with the N-terminal region, with a focus point on H69 and K71 which are part of a small alpha helix (Fig 2B). Reduced EVI1-CTBP2 interaction was predicted with AlphaFold when adjacent residue V72 in CTBP2 was mutated into an arginine (V72R) (Fig 2B). Indeed using MAPPIT, we determined that in CTBP2 the valine residue at position 72 (V72) was important for binding EVI1, whereas alanine at position 58 (A58) was not (Fig 2E). In summary, with its PLDLS site, EVI1 recognises a specific HEKV binding region in CTBP2.

The importance of the PLDLS motif in EVI1-mediated transformation was studied in mouse bone marrow progenitors in vitro. Transduction with a retroviral construct carrying wild type Evi1 formed colonies of cells that can be replated indefinitely (Fig 2F). Serial replating capacity of mouse bone marrow cells transduced with mouse Evi1 lacking the PLDLS domain was severely reduced, demonstrating the importance of this specific domain and its interactor CTBP2 for EVI1-mediated transformation of bone marrow progenitors (Fig 2F).

### Overexpression of a PLDLS-competitor causes loss of EVI1 to CTBP2 interaction

Next, we wondered whether it is possible to interfere between EVI1 and CTBP2 using a competing PLDLS-containing construct. First, we determined the minimal region around PLDLS required for a competitor peptide to bind CTBP2. An EVI1-PLDLS construct with 10-15 amino acids (AAs) at either side of PLDLS fully induced the STAT3-driven luciferase activity in the MAPPIT assay with CTBP2 as prey protein (Fig. S3B). With AlphaFold, a 1x PLDLS repeat was predicted to bind CTBP2 at the same site where full-length EVI1 binds (Fig 3A). We next designed a construct with 4 repeats of the PLDLS motif including 15 flanking AAs from EVI1 on either side of each PLDLS site (Fig 3B). The 4x PLDLS construct was able to fully outcompete the EVI1/CTBP2 interaction in the MAPPIT assay, whereas a similar 4x PLASS control did not affect EVI1/CTBP2 interaction (Fig 3C). In line with this, EVI1-IP/CTBP2-WB demonstrated loss of CTBP2 binding to EVI1 in the presence of 4x PLDLS, whereas 4x PLASS construct did not affect EVI1-CTBP2 interaction in MUTZ3 cells (Fig 3D, Fig. S3D). The interaction between EVI1 and CTBP1 was also specifically outcompeted by 4x PLDLS, indicating that both proteins can bind EVI1 via the same motif (Fig 3D). Upon transduction of MUTZ3 cells with 4x PLDLS, CTBP2 binding to chromatin was globally lost, whereas 4x PLASS control did not affect CTBP2 interaction with chromatin (Fig 3E,Fig 3F,Fig. S4A). This shows that in 3q26/MECOM-rearranged leukemia cells the effect of overexpression of the 4x PLDLS inhibitor on CTBP2 binding to chromatin is similar to EVI1 knock-down (Fig. 1E). In addition, while downregulation of CTBP2 binding to chromatin was global, it was even stronger in CTBP2 peaks that overlap with an EVI1 binding site in ChIP-seq for MUTZ3 (Fig. S4B). We next studied the effect of 4x PLDLS overexpression on the transcriptome of MUTZ3 cells. We identified 162 significantly differentially expressed genes (126 upregulated and 36 downregulated), of which 70% of up-regulated and 50% of down-regulated genes contained at least 1 CTBP2 binding site in their locus (Fig. S4C). 29 of the top-30 CTBP2 peaks that are annotated to 4x PLDLS target genes are upregulated genes in RNAseq, in line with a repressor function of EVI1-CTBP2 (Fig. S4D).

**Figure 3.**
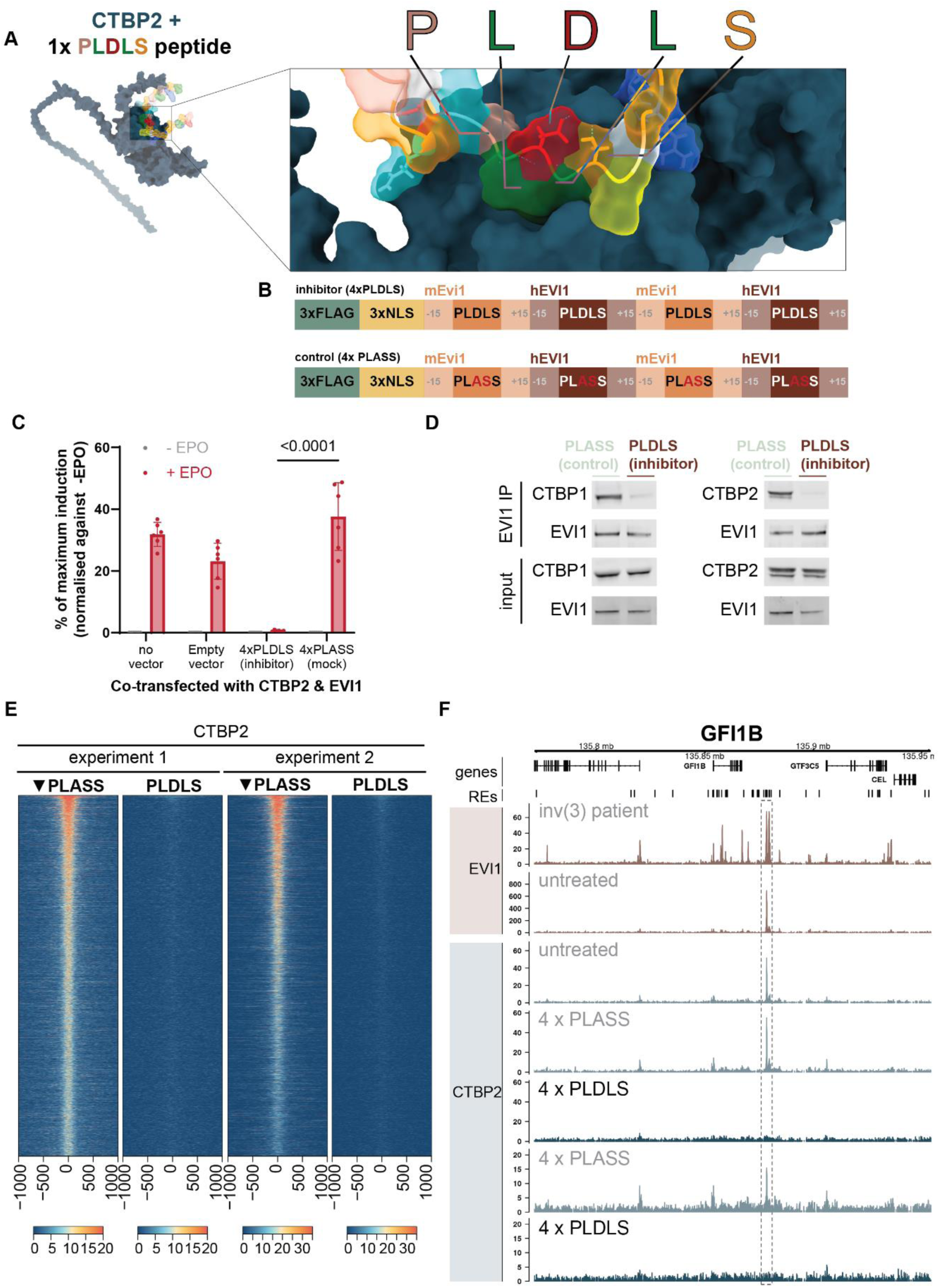
Disruption of CTBP2 binding to EVI1 with PLDLS inhibitor. A. AlphaFold prediction of the interaction of the PLDLS inhibitor with CTBP2 visualized with ChimeraX. The PLDLS residues are shown in different colours according to their biochemical properties and CTBP2 is depicted in dark blue. B. Diagram outlining the 4xPLDLS inhibitor and 4xPLASS control constructs. Each repeat consists of two human (hEVI1) and two mouse (mEvi1) PLDLS surrounding region (±15 AAs); NLS = nuclear localisation signal. C. MAPPIT assay in HEK293T cells to measure EVI1-CTBP2 interaction as in figure 2 with 4xPLDLS inhibitor or 4xPLASS control. Reporter induction is normalised to –EPO (not induced) and maximum induction with RNF as positive control. Significance was determined with two-way ANOVA (comparison PLDLS vs PLASS + EPO shown). D. Western blot for CTBP1 and CTBP2 following IP for EVI1 in MUTZ3 cells with dox-induced 4x PLASS or 4xPLDLS expression. E. Heatmap of ChIP-seq data for CTBP2 in MUTZ3 cells following retroviral transduction with either 4x PLASS (control) or 4x PLDLS (inhibitor). Bona fide CTBP2 binding sites were determined by overlapping the two control (PLASS) tracks (5984 peaks). Tracks were ranked combined enrichment at these overlapping peaks, which was calculated by multiplying the normalized peak enrichment of the control tracks with each other. Peaks were ranked on this combined enrichment. F. Representative plot showing the effect of 4x PLDLS overexpression on the locus of GFI1B. (RE = regulatory element associated with expression of GFI1B)

### The PLDLS competitor abolishes leukemic potential of EVI1-transformed mouse bone marrow cells

We next investigated whether interference between EVI1 and CTBP2 using by 4x-PLDLS affects Evi1-driven transformation in mouse bone marrow cells (Fig 4A). Colony assays demonstrated that the replating ability of Evi1-transformed bone marrow was fully abolished when these cells were transduced with a 4x PLDLS competitive inhibitor, whereas the 4x PLASS mock inhibitor had no effect (Fig 4B,Supplementary Table 1). In E2a-Pbx transformed bone marrow, the effect of 4x PLDLS on colony formation was significantly smaller than in Evi1-transformed bone marrow (Fig 4B). In the presence of 4x PLDLS, but not 4x PLASS, we observed strong neutrophilic differentiation of the Evi1-transformed bone marrow cells determined by flow-cytometric analysis (Gr1+/CD11b+) (Fig 4C) and by May–Grünwald Giemsa staining of cytospins (Fig 4D). Those experiments indicate that the differentiation arrest of myeloid progenitors caused by Evi1 overexpression depends on the interaction with CTBP1/2. Together, these results indicate that Evi1-transformed cells are unable to sustain their leukemic potential in presence of the PLDLS inhibitor.

**Figure 4.**
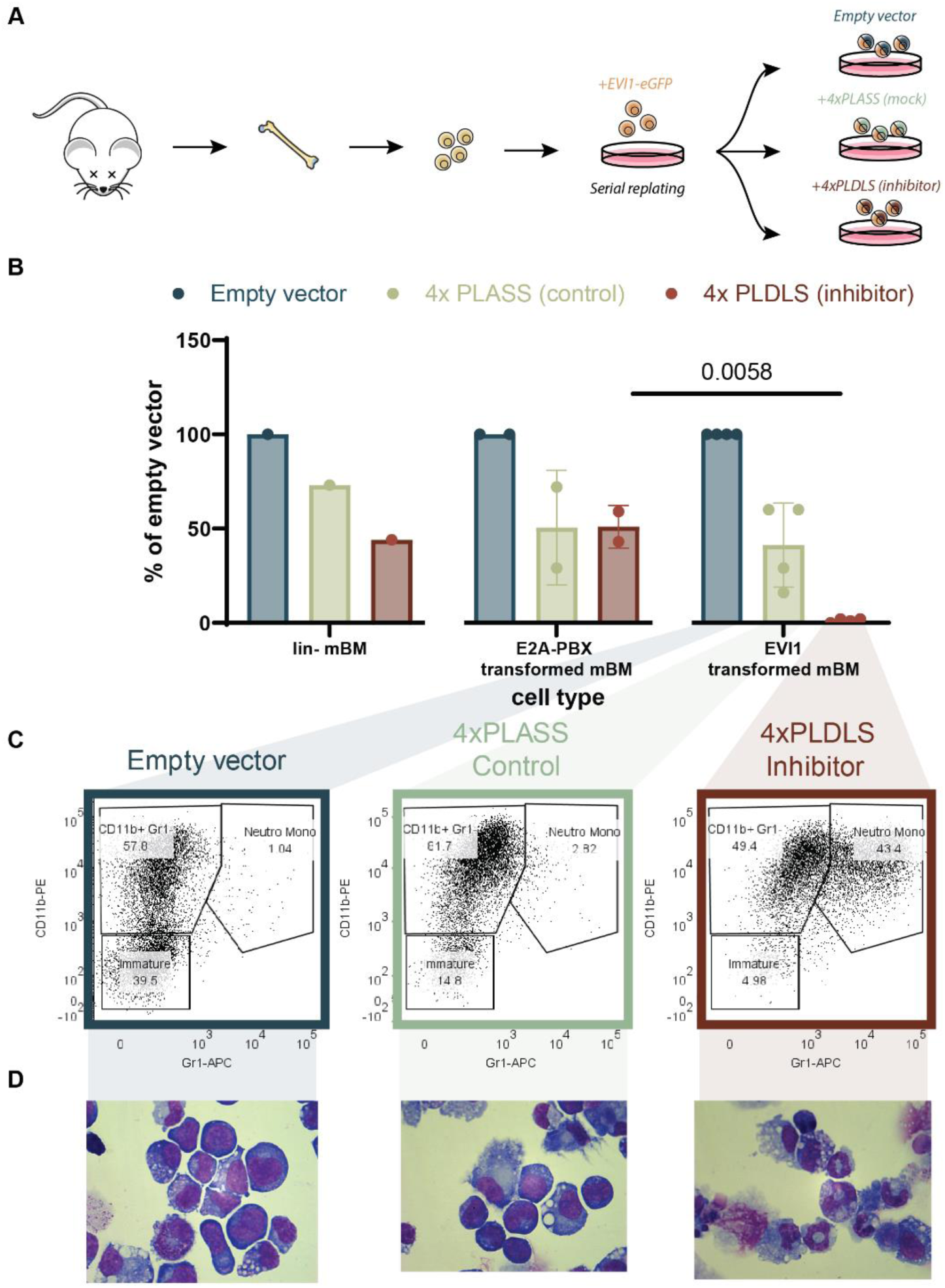
Inhibition of EVI1-CTBP2 in transformed mouse bone marrow abolishes leukemic potential. A. Diagram illustrating the procedure for functionally testing inhibitors of the EVI1-CTBP interaction. For all experiments on EVI1-transformed bone marrow shown, bone marrow of 8- to 10-week-old mice were collected and transduced with a retroviral EVI1-eGFP construct. The immortalised cells acquired after 4 rounds of serial replating were transduced with retroviral vectors containing no insert (empty vector), a 4x PLASS control, or 4x PLDLS (inhibitor). B. Colony formation of lineage-negative (Lin-) mouse bone marrow cells (left panel), Lin-cells immortalised with E2A-PBX (middle), or with EVI1 (right panel). Prior to plating, cells were transfected with empty vector, PLASS (control) or PLDLS (inhibitor) and selected with puromycin. All experiments were performed in triplicate; each dot represents the average of technical triplicates of an independent experiment (n=1-4). Adjusted P-value of two-way ANOVA with Šídák’s multiple comparisons test shown if adjusted P-value < 0.05. Between-group comparison shown on graph; within-group test results are included in Supplementary Table 1. C. Flow -cytometric analysis of Lin-mouse bone marrow transduced with EVI1, and with empty vector, 4xPLASS or 4xPLDLS, stained with CD11b-PE and Gr1-APC (single cell gate shown) (Day 7; cultured in media with cytokines including GM-CSF). D. May-Grünwald Giemsa staining of cytospins made from Lin-mouse bone marrow transduced with EVI1, together with empty vector, 4xPLASS or 4xPLDLS (Day 7; cultured in media with cytokines including GM-CSF)((same experiment as 4C).

### The PLDLS inhibitor blocks in vivo outgrowth of human EVI1-transformed AML cells

Next, we investigated the effect of the 4x PLDLS inhibitor on EVI1-transformed human leukemia cells. Overexpression of 4x PLDLS strongly reduced in vitro colony formation of the EVI1-positive 3q26-rearranged AML cell lines MUTZ3 and SB1690CB, compared to colony formation in the presence of 4x PLASS (Fig 5A,Supplementary Table 1). The effect was much stronger than in EVI1-negative cell line HL60 (Fig 5A). To measure the direct competition between cells expressing either 4x PLASS or 4x PLDLS, we transduced SB1690CB cells with either PLDLS-or PLASS-containing vectors. To enable cell sorting and tracking over time, each vector contained a fluorescent read-out (mCherry or Emerald). At day 0, cells were sorted and mixed in approximately 1:1 ratio of PLDLS- and PLASS-containing fractions. Two replicate cell line mixtures were generated: PLDLS-mCherry/PLASS-Emerald and PLDLS-Emerald/PLASS-mCherry. PLASS-containing SB1690CB cells always outcompeted PLDLS-transduced SB1690CB cells in vitro (Fig 5B, Fig 5C). The effect of the 4x PLDLS inhibitor on the outgrowth of EVI1 expressing AML cells in vivo was studied in two xenotransplant models. First, the mCherry+ and Emerald+ mixtures of SB1690CB cell were injected into 6 mice in a 1:1 ratio (Fig 5D). Flow-cytometric analysis at input showed the equal distribution of mCherry+ and Emerald+ cells at the start of the experiment, whereas after 10 weeks virtually only PLASS-containing cells were detectable (Fig 5D, Fig S5A, Fig S5B). In a second in vivo model, we transplanted human MUTZ3 cells into NSG mice with surgically implanted human bone marrow scaffolds (Fig 5F) (31, 40). The MUTZ3 cells transplanted into these mice contained a CMV-promoter-driven luciferase construct, enabling live measurement of graft size over time. Outgrowth of MUTZ3-luciferase cells with 4x PLDLS inhibitor was strongly reduced compared to 4x PLASS control cells (Fig 5G, Fig 5H, Fig S5C). Tumours that were detectable at the implanted scaffolds in mice transplanted with 4x PLDLS-containing cells were smaller than those in scaffolds with 4x PLASS-containing MUTZ3 cells (Fig 5H). Thus, human EVI1-transformed AML cells depend on the PLDLS-dependent interaction between CTBP and EVI1, which can be targeted to reduce the growth of this leukemia in vivo.

**Figure 5.**
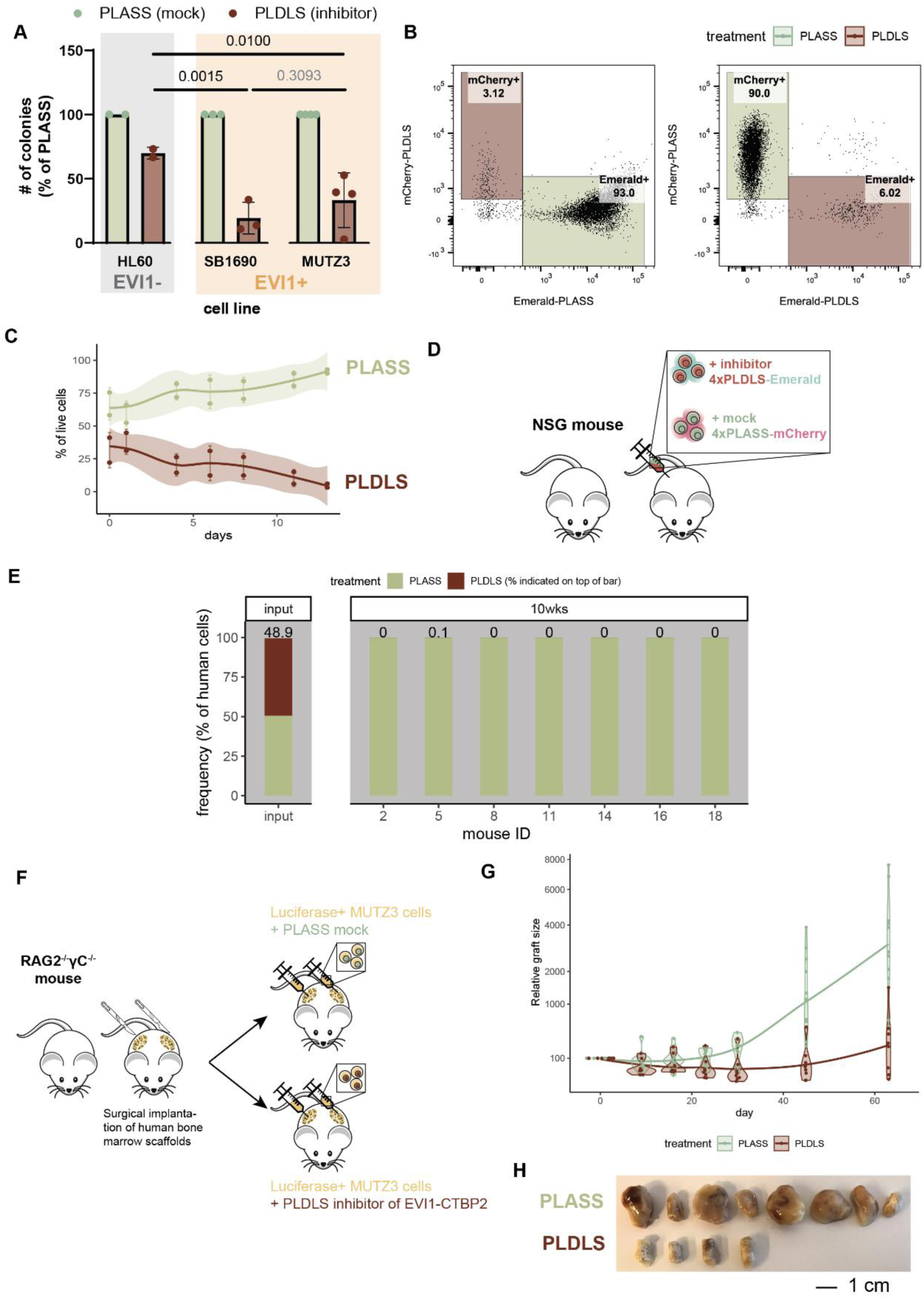
In vivo growth inhibition of EVI1-transformed human leukemia cells using EVI1-CTBP2 inhibitor. A. Colony formation of human EVI1-(HL-60) or EVI1+ (SB1690 or MUTZ3) cell lines, which are transduced with a retroviral vector with either 4x PLDLS or 4x PLASS. All experiments were performed in triplicate, n = 2-4 experiments per cell line (2-way ANOVA with multiple testing correction). Between-group test results shown on graph; within-group test results shown in Supplementary Table 1. B. SB1690 cells were transduced with lentiviral constructs, sorted and mixed to 1:1 ratio Emerald: mCherry on day 0. Flow-cytometric analysis at Day 13 is shown. The left panel shows a mix of 4xPLDLS-mCherry and 4xPLASS-Emerald. The right panel displays a mix of 4xPLDLS-Emerald and 4xPLASS-mCherry. C. Competition between 4xPLASS and 4xPLDLS in SB1690CB over time. Quantification of Emerald and mCherry gates from panel 5B, grouped according to whether they contain inhibitor (PLDLS) or control (PLASS) sequence. D. Diagram illustrating competition assay in NSG mice using SB1690CB cells. Before transplant, SB1690 cells were separately transduced with PLDLS-Emerald or PLASS-mCherry and mixed in a 1:1 ratio. Aliquots of the cell mixture were injected into seven mice and after 10 weeks bone marrow cells were isolated and analysed with flow cytometry. E. Flow-cytometric analysis of input (at the start of the experiment) and 7 mice transplanted (at the end of the experiment (week 10)) with SB1690. Percentage of 4xPLDLS-positive cells is indicated above each bar. F. Diagram illustrating the procedure to engraft human MUTZ3 cell line into NSG mice. Before transplantation, human bone marrow scaffolds are surgically implanted into NSG mice. Mice are then injected with either MUTZ3-Luciferase cells +4x PLDLS inhibitor or MUTZ3-Luciferase cells + 4xPLASS mock. Upon injection with luciferin, mice can be imaged in a luminescence scanner to monitor outgrowth of MUTZ3 cells (Fig. S5A) G. Tumour area of mice who received either a transplant of MUTZ3-Luciferase cells + 4xPLDLS inhibitor or MUTZ3-Luciferase cells + 4xPLASS mock (n = 4 mice with 2 scaffolds each for PLDLS; and n = 6 mice with two scaffolds each for PLASS). H. Scaffolds of mice with engraftment of 4xPLASS or 4xPLDLS

## Discussion

AML patients with overexpression of EVI1 caused by inv(3) or t(3;3) have an extremely poor prognosis and are frequently refractory to current treatments (1). Consequently, there is a major need for the development of new ways to treat these patients. We have previously demonstrated that the survival and growth of those leukemia cells heavily depend on EVI1 (3). Here, we show that EVI1 interacts with CTBP1/2, that the interaction is essential for leukemic transformation and that it can be targeted. The interaction between EVI1 and CTBP1/2 has previously been reported (10, 38), and also recently in hematopoietic cell lines (41), but its importance in AML development was not shown. We found that in AML cells CTBP2 predominantly interacts with EVI1 via a conserved PLDLS motif, which was previously described to mediate interaction with CTBP1 in rat fibroblasts (38). We demonstrate here that without this PLDLS motif or following overexpression of a PLDLS competitor construct, transformation by EVI1 in 3q26/MECOM-rearranged AML is abolished.

A key question that remains is whether inhibition via a PLDLS domain (which is present in more proteins than EVI1) specifically targets EVI1, and whether EVI1- expressing cells are uniquely sensitive to this type of inhibition. CTBP2 interacts with ZEB2, and using the MAPPIT assay we showed that the PLDLS inhibitor affects this interaction as well (Fig. S3C). Likewise, CTBP2 can interact with E1A, AREB6 or FOG (42). In addition, it has been shown that E1A and CTBP1 can be outcompeted with a E1A mimicking peptide as well (43). However, we found global loss of CTBP2 binding to chromatin upon EVI1 knockdown or when the 4x PLDLS inhibitor was overexpressed. This indicates that EVI1 is critical for CTBP2 binding in 3q26/MECOM-rearranged AML cells, and that ZEB2 or other PLDLS containing proteins present in the cells are unable to functionally replace EVI1 in tethering CTBP2 to chromatin. In addition, we demonstrate less sensitivity to PLDLS inhibition in HL60 cells, mouse bone marrow progenitors and E2A-PBX transformed mouse bone marrow, none of which express EVI1. Thus, even though other proteins might be affected by inhibition by 4x PLDLS, the fact that EVI1-expressing cells are more sensitive to the effect of PLDLS inhibition suggests that there is a therapeutic window for targeting this interaction in AML patients.

It is unclear what the biological role of CTBP1/2 is in the transformation process. Perhaps their only function is to serve as scaffold proteins that bring other critical proteins to various specialized complexes. While CTBP1 and CTBP2 are highly enriched in MS-IP for EVI1 and important for EVI1 function, many other proteins interact with EVI1 and may also play a role in transformation. For instance, we identified HDAC1/2, RCOR1, RUNX1 and EHMT1/2 among ∼400 significant proteins in our IP-MS experiments which are known interactors of CTBP1/2. In addition, there might also be important interactions of EVI1 that are not mediated by CTBP1/2. The importance of CTBP-dependent or CTBP-independent EVI1 interactors in leukemic transformation should be further investigated using a genetic knock-out screen. It is also possible that there is an additional role for CTBPs in modulating chromatin bound by EVI1. CTBP1 and CTBP2 are highly structured proteins that contain two large oxireductase enzymatic domains (44). Both are annotated as catalysts of a NAD+ to NADPH conversion in the methionine salvage pathway, even though this has been studied mostly for CTBP1 (45–47). NAD binding has been reported to be important for homo- or heterodimerization of CTBPs with each other (48). It is unclear what substrate CTBP1/2 catalyze while in complex with EVI1, and whether the product of this chemical reaction is limiting for any other metabolic reaction taking place at chromatin. It would be interesting to investigate the effect of mutating enzymatic domains in CTBP1/2 on transformation by EVI1. Metabolic profiling of nuclei, perhaps incorporating spatial information in the presence of our PLDLS inhibitor, could reveal whether CTBP1/2 has any local effects on chromatin that might be important for gene regulation.

Taken together, in this proof-of-concept study we show that interference with protein-protein interfaces of transcription factors such as EVI1 and their cofactors can inhibit tumor growth and should be exploited for therapeutic purposes. Since EVI1 overexpression has also been reported in other tumor types, such as ovarian or breast cancer, our findings may not be restricted to 3q26/MECOM-rearranged AMLs. Whether PLDLS-to-CTBP interaction is essential in those tumors needs to be investigated. Generation of chemically optimised EVI1-derived PLDLS peptides might lead to the development of stable small molecules that interfere with EVI1 and CTBP1/2 complex formation. Such compounds could ultimately be tested in a clinical setting, which may provide a new treatment option for this highly aggressive form of AML caused by overexpression of EVI1.

## Materials and Methods

### Data availability

ChIP-seq data is made available on GEO under accession number GSE236010. Raw mass spectrometry data and data for protein identification and quantification is submitted to the ProteomeXchange Consortium via the PRIDE partner repository with the data identifier PXD043333. All other data used for generating figures presented here (AlphaFold predicted structures, flow cytometry frequencies, luciferase output files and mass spec clustering and quantification) are available at Zenodo under accession 10.5281/zenodo.8354861. Custom scripts for AlphaFold quantifications and ChIP-seq correlation plots are available on Github (https://github.com/dorienpastoors/EVI1_CTBP2_PLDLSinhibitor). An overview of all data deposited associated with this manuscript is available in Supplementary Table 1.

### Cell lines

MUTZ3 (DSMZ, cat ACC295) cells were cultured in Alpha-mem medium (Gibco cat 2571038) supplemented with 20% supernatant of urinary bladder carcinoma cell line 5637 (DSMZ, cat ACC35) (17), and 20% FCS (Sigma-Aldrich, cat F7524-500ml). SB1690CB cells were cultured in RPMI (GIBCO, cat 21875-034), 10% supernatant 5637, 10% FCS and 10 ng/ml human IL3 (Peprotech, cat no 200-03). MOLM1, HNT34, HL60 (DSMZ, cat ACC 720,600 and 3) were cultured in RPMI and 10% FCS. 293T (DSMZ, cat ACC635) were cultured in DMEM (GIBCO, cat 2430-025) and 10% FCS. All medium used was supplemented with 50U/ml penicillin and 50 U/ml streptomycin (Gibco, cat 15140122). All cell lines were routinely confirmed to be mycoplasma free by using the MycoAlert Mycoplasma Detection kit (Lonza, cat no LT07-318). SB1690CB was a gift of Stefan Meyer (18).

### Generation of viral supernatants

Retroviral supernatants were prepared by co-transfecting 293T cells (DSMZ, cat ACC635) using pCL ECO (Addgene, cat 12371) for mouse cells or pCL ampho for human cells and a retroviral vector. Retrovirus was harvested at 48 hours post transfection. Lentiviruses were produced by co-transfecting 293T cells using psPAX2 (Addgene, cat 12260), pMD2.G (Addgene, 12259) and a lentiviral vector. Lentiviral supernatants were harvested at 72 hours post transfection. All transfections were performed using Fugene-6 transfection reagent according manufacturers protocol (Promega, cat E2691).

### Overexpression EVI1 / PLDLS EVI1 mutant in murine HCS and colony forming assays

8- to 10-week-old female C57bl/6 mice (Jackson Laboratory) were euthanized and bone marrow was isolated from their femurs and tibias. Erythrocytes were lysed by resuspending total bone marrow in ACK-lysing solution (Gibco, cat A10492-01). Mononucleated cells were enriched for hematopoietic progenitor stem cells following manufacturers protocol (BD, cat 558451). After isolation, cells were cultured overnight in Cellgro SCGM medium (Cellgenix, cat 2001) supplemented with 1:5 Angiopoietin, 10 ng/ml SCF, 10 ng/ml FGF, 20 ng/ml TPO and 20 ng/ml IGF (PeproTech) (19). Cells were transduced using pMYs-EVI1 (a kind gift of Mineo Kurokawa), pMYs-ΔEVI1 (single mutation in PLDLS motif) or an empty vector control by Retronectin transduction. In short: 3.5 cm non-coated cell culture dishes (Falcon, cat 351008) were coated using 12 µg Retronectin (Takara, cat T100B). Coated dishes then were pre-incubated with 1 ml viral supernatant for 4-5 hours at 37oC. After removing the virus, 1-2x106 HSC’s from an overnight culture were added to each dish. Cells were cultured overnight. This procedure was repeated for an additional 24 hours. After transduction, cells were plated in MethoCult (Stemcell, cat M3231) containing 10 ng/ml IL3, IL6, GM-CSF and SCF (all from Peprotech) at a concentration of 3x104 cells per/ml. Colonies were counted and replated every 7 days. Material harvested from serial replatings was used in expression analysis for EVI1 by RT-qPCR. After 4 rounds of replating, EVI1 overexpressing cells were transferred to RPMI medium (GIBCO, cat 21875-034) containing 10 ng/ml IL3, IL6, GM-CSF and SCF (Peprotech) to establish a multiclonal cell line overexpressing EVI1. Data visualization and statistical testing was performed in GraphPad Prism (v 9.3.1).

### RNA-sequencing

MUTZ3 cells were transduced with retroviral supernatants encoding either p50MX-4x empty vector-Zeocin, p50MX-4x PLASS-Zeocin or p50MX-4xPLDLS-Zeocin. RNA was isolated at 72HRS of Zeocin selection. RNA quality was determined and 1 ug RNA was used in library preparation using TruSeq RNA library preparation kit (Illumina, cat no RS-122-2001). Library preparation was performed according manufacturers protocol. DNA was sequenced on a Illumina 2000-2500 platform, 75 cycles PE.

### RT-PCR of human and mouse EVI1

A minimum of 1x106 cells were resuspended in 1ml Trizol (Life Technologies, cat 15596018) and RNA was isolated following manufactures protocol. 1 µg RNA was used in reverse transcription using Superscript II reverse transcriptase (Life Technologies, cat 18064-014). Human or mouse EVI1 PCR was carried out using specific primer sets (see Supplementary Table 1) using an ABI7500 real-time PCR cycler. Expression was normalized using mouse Hprt or human PBGD.

### Western blot analysis

Cells were lysed in Sarin buffer (20 mM Tris-HCl, 138 mM NaCl, 10 mM EDTA, 50 mM NaF, 1% Triton, 10% glycerol) which was supplemented with protease inhibitors SigmaFast, Na3VO4 and reducing agent DTT (all chemicals purchased from Sigma-Aldrich). Protein content was measured by Pierce BCA protein Assay (Thermo scientific, cat 23227). 40 µg protein was loaded on a 4-12% Bis-Tris polyacrylamide gel (Thermofisher cat NP0321) and ran in a Mini Gel Tank (Thermofisher) in 1X MOPS buffer (Thermofisher cat NP001) Proteins were semi-dry blotted onto a 0.2 µM nitrocellulose membrane (Sigma cat GE10600001) and protein levels were detected by specific antibodies directed against: Human EVI1 (Cell signaling, cat 2265), CTBP1 (BD, cat 612042), CTBP2 612044 (BD, 612044), V5 tag (Life Technologies, cat R96025), FLAG tag (Sigma-Aldrich, cat F3165), β-actin (Sigma-Aldrich, cat no A5441). HA tag (Santa Cruz cat no Sc-805). Proteins were visualized using the Odyssey infrared imaging system (LI-COR Biosciences).

### Flow Cytometry

Flow-cytometric analysis of mouse bone marrow cells or MUTZ3 cells was done with specific antibody stainings using mouse CD11B-APC (BD, cat 553311) and mouse Gr-1 FITC (BioLegend, cat 108406). Cells were analyzed on a BD LSRII flow cytometer (BD Bioscience). Data was analysed with FlowJo (dotplots, and frequency tables). All other graphs were made in ggplot.

### ChIP-seq

Chromatin immunoprecipitation was performed in MUTZ3, MOLM1, HNT34, and SB cells using specific antibodies against human EVI1 (Cell Signaling, cat 2593) and CTBP2 (BD, 612044). Cells were crosslinked for 45 minutes using 0.5 µM DSG (ThermoFisher scientific, cat 20593). Cells were pelleted, resuspended in PBS and crosslinked for 10 minutes using 1% formaldehyde (Sigma-Aldrich, cat F8775-500ml). Formaldehyde crosslinking was quenched by adding 1.25M glycin 5 minutes at RT. Cells were washed 3 time using cells lysis buffer containing 10 mM Tris-HCL pH 7.5, 10 mM NaCl, 3mM MgCl, 0.5% NP40. After last wash, cells were resuspended in sonication lysis buffer with, 0.8% SDS, 160 mM NaCl, 10mM Tris-HCL pH 7.5, 10mM NaCl, 3 mM MgCL, 1 mM CaCl2 and 4% NP40. Lysates were sonicated on a biorupter pico (Diagenode). Chromatin was diluted 4-5x using IP dilution buffer containing 1.1% Triton X-100, 0.01% SDS, 167 mM NaCl, 16.7 mM Tris-HCL pH 8.0 and 1.2 mM. 2% sample was saved as an input control. 5 µg antibody was added and chromatin mixed O/N at 4oC. The following day, 30 µl protein G beads (ThermoFisher scientific, cat 1004D) was added to the samples and mixed for an additional 3 HRS. Washing of the beads and DNA elution was performed following the standard ChIP protocol from Upstate. (All chemicals in buffers are purchased at Sigma-Aldrich). Libraries for sequencing were generated by using either the Truseq ChIP (BIOO Scientific / Perkin Elmer, cat nova-5143-01) or MicroPlex Library v3 (Diagenode, cat C05010001) preparation kit according manufacturers protocol. Samples were sequenced single-end (1 × 50 bp) on the HiSeq 2500 platform (Illumina) or paired-end (2 × 100 bp) on the Novaseq 6000 platform (Illumina).

### MAPPIT assay

HEK293T cells were seeded at 0.35e6 cells/well in a 6 well plate. The next day, cells were transfected with 1 μg pXP2d2-rPAP1-luciferase (STAT3-responsive luciferase reporter), 1μg pCEL (the modified EPO receptor; bait) and 1μg pMG2 (the gp130 fusion needed to activate STAT3 at the modified receptor; prey) in 250 μl DMEM -/-/-medium containing 9μL lipofectamine-2000 (Invitrogen #11668500). For each condition, a positive control (MG2-RNF) was also transfected to which data is normalised and which represents maximum reporter induction. DNA, media and lipofectamine are incubated for 5 minutes at room temperature and added dropwise to cells. After 48 hrs of transfection, each well was trypsinized and divided over 4 wells in a 96-well plate in a total volume of 100μL, and cells were incubated o/n with EPO (4U/μL) to induce JAK/STAT signaling for 24 hrs. Luciferase activity was measured with addition of 50 μL Steady-Glo (Promega cat #E2510), incubated for 20 minutes, and luminescence was measured with a Victor-X4 (Perkin-Elmer). For each experiment, replicates were collapsed when baseline (-EPO) was subtracted and data was normalised to % induction of the positive control (RNF) (-EPO was 0). Experiments from independent dates were combined and significance tested in a 2-way ANOVA (or mixed model in the case of missing values). P-values reported are adjusted for multiple testing. Data visualization and statistical testing was performed in GraphPad Prism (v 9.3.1).

### Mass spectrometry

#### Nuclear protein extractions for mass spectrometry

Nuclear extractions were performed as previously described (20). Briefly, cells were spun down (500g 5 min at 4 °C) and washed twice in cold PBS. Pellet volume was determined using a standard and cells were resuspended in 5 volumes cold buffer A (10mM HEPES KOH pH 7.9, 1.5mM MgCl2, 10mM KCl) and incubated for 10 minutes on ice, and centrifuged (400g 5 minutes 4 °C). Cell volume was determined again and 2 volumes cold buffer A++ (Buffer A + cOmplete protease inhibitors (Roche, cat 04693159001) and 0.15% NP40). Cell suspension was transferred to a pre-cooled, buffer A calibrated dounce homogenizer. 4x 10 strokes with a type B pestle (tight) were perfomed, and the douncer was left to cool for 30 seconds after each 10th stroke. The suspension spun down for 15 mins at 3900 RPM (large centrifuge) and the cytoplasmic extract (the supernatant) was not used in this study. The pellet was washed in in 2-3 mL cold PBS by flicking the tube and spun down again at 3900 RPM for 5 minutes. The pellet size was determined with the standard and 2 volumes of cold Buffer C+++ (420mM NacL, 20mM HEPES KOH pH 7.9, 20% v/v glycerol, 2 mM MgCl2, 0.2mM EDTA, 0.1% NP40, cOmplete protease inhibitors (See buffer A++) and 0.5mM DTT) were added and vigourously resuspended (not vortexed). The lysate was rotated for 1 hr at 4 °C and spun down (Table-top centrifuge, 14000RPM 30 minutes at 4 °C) and the supernatant (nuclear extracts) were collected, snap-frozen and stored at −80 °C until further use.

#### Immunoprecipitation of EVI1 for mass spectrometry and western blot

Concentration of nuclear protein extracts was determined using the Bradford assay (ThermoFisher, cat #23238). All MS-IP experiments were done in triplicate, with cell lysates obtained on separate dates and the IP performed simultaneously. All steps were performed on ice and all centrifuge steps at 4 °C. For each IP on nuclear lysate, 1 mg nuclear lysate was adjusted to 500 uL in Protein Incubation Buffer (PIB+++; 150mM NaCl, 50 mM Tris pH 8.0, cOmplete protease inhibitor (Roche, cat 04693159001), 0.25% NP-40 and 1 mM DTT). For WB-IP on total cell lysate, the IP and all washing steps were performed in the protein extraction Sarin buffer (see Western blot). 30 uL ProtG magnetic Dynabeads (ThermoFisher, cat 1003D) were washed 3 times in 1 mL PIB+++ or Sarin buffer. Lysates were pre-cleared by mixing head-over-head for 2 hrs at 4 °C. For input samples for western blot, 80ug protein lysate was heated at 95 °C for 5 mins in 40 uL 1x Protein loading buffer (sample buffer for SDS-Page gels). For the IP, 50 uL ProtG Dynabeads (see above) were washed in 3x1mL PIB+++ or Sarin and added to the pre-cleared lysates. 5 ug primary antibody was added (IgG control: Cell Signaling Rabbit (DA1E) mAb IgG XP® Isotype Control #3900 Lot5; EVI1: Cell Signaling C50E12 Anti-EVI1 Rabbit IgG #2593S Lot3, CTP2: BD, #612044) and samples were mixed head-over-head overnight at 4 °C. Samples were washed 5x in 1mL cold PBS + 0.1% NP40 or Sarin buffer and processed for mass spectrometry. For western blot samples, proteins were eluted from the beads with 30 uL 1.5x Protein loading buffer and heated 10 minutes to 95 °C while shaking and loaded onto a western blot gel (see Western blot section).

#### Mass spectrometry analysis

Nanoflow LC-MS/MS was performed on an EASY-nLC system (Thermo) coupled to a Orbitrap Fusion Lumos Tribrid mass spectrometer or an Orbitrap Eclipse Tribrid mass spectrometer (both Thermo), operating in positive mode and equipped with a nanospray source. Peptide mixtures were trapped on a ReproSil C18 reversed phase column (Dr Maisch GmbH; column dimensions 1.5 cm × 100 µm, packed in-house) at a flow rate of 8 µl/min. Peptide separation was performed on ReproSil C18 reversed phase column (Dr Maisch GmbH; column dimensions 15 cm × 50 µm, packed in-house) using a linear gradient from 0 to 80% B (A = 0.1% FA; B = 80% (v/v) AcN, 0.1 % FA) in 120 min and at a constant flow rate of 250 nl/min. The column eluent was directly sprayed into the ESI source of the mass spectrometer.

For data dependent acquisition (DDA): All mass spectra were acquired in profile mode. The resolution in MS1 mode was set to 120,000 (AGC target: 4E5), the m/z range 350-1400. Fragmentation of precursors was performed in 2 s cycle time data-dependent mode by HCD with a precursor window of 1.6 m/z and a normalized collision energy of 30.0; MS2 spectra were recorded in the orbitrap at 30,000 resolution. Singly charged precursors were excluded from fragmentation and the dynamic exclusion was set to 60 seconds.

#### Quantification

DDA raw data files were analyzed using the MaxQuant software suite (version 2.2.0.0 (21)) for identification and relative quantification of proteins. ‘Match between runs’ was disabled and a false discovery rate (FDR) of 0.01 for proteins and peptides and a minimum peptide length of 6 amino acids were required. The Andromeda search engine was used to search the MS/MS spectra against the Homo sapiens Uniprot database (version May 2022) concatenated with the reversed versions of all sequences and a contaminant database listing typical background proteins. A maximum of two missed cleavages were allowed. MS/MS spectra were analyzed using MaxQuant’s default settings for Orbitrap and ion trap spectra. The maximum precursor ion charge state used for searching was 7 and the enzyme specificity was set to trypsin. Further modifications were cysteine carbamidomethylation (fixed) as well as methionine oxidation. The minimum number of peptides for positive protein identification was set to 2. The minimum number of razor and unique peptides set to 1. Only unique and razor non-modified, methionine oxidized and protein N-terminal acetylated peptides were used for protein quantitation. The minimal score for modified peptides was set to 40 (default value).

### Data analysis

#### Data visualisation

Analysis have been conducted using R version 4.2.2 and visualized with ggplot2 unless otherwise specified.

#### Statistical analysis

Statistical analysis is specified in the corresponding method section of the analysis type concerned. In general, for colony assays or plate-based data such as MAPPIT, GraphPad prism is used with two-way ANOVA (with appropriate multiple-testing corrections). For other data types, specialized statistical software is used (DEseq2 for RNA-seq, DiffBind for ChIP-seq data, Perseus for mass-spectrometry data).

#### Differential enrichment analysis Mass Spectrometry

Perseus (version 2.0.3.1) was used for differential enrichment analysis. Log2 LFQ intensities from MaxQuant were loaded into Perseus. Common contaminants were excluded and rows were filtered that contained values in at least 3 samples within a group. Missing values were replaced by a value below the lowest detected protein. DE analysis was performed with the following parameters: Program: EdgeR; Test method: Likelihood ratio test; Normalization method EdgeR: TMM). Data for volcano plots was generated with a T-test on the normalized data (Parameters: Side: Both; Number of randomizations: 250; Preserve grouping in randomisations: None; FDR threshold: 0.05). The data were exported from Perseus and visualized in R using ggplot2. Biogrid database was used to identify previously identified proteins (version 4.4)(22). Complex enrichment was performed with EnrichR with the CORUM database (23). For the network analysis, a graph of proteins from the Perseus analysis (nodes) with connections (edges) representing their co-occurrence in the CORUM complexes was constructed using the Fruchterman-Reingold layout algorithm. The edges were weighted on the basis of the cosine distance between proteins in terms of how frequently they co-occur in the same complex. Clusters or communities of proteins with shared complexes were detected using the Louvain algorithm with default resolution. These analyses were conducted with the tidygraph and igraph R packages, and visualized with ggraph. For the MS experiment in NFS78 (Fig S1D), data from three independent experiments were pooled. All control samples (NFS78 untransduced, Bir-A only) were compared to experimental samples (EVI1-Biotag +BirA expressing clones). Before importing them into Perseus, Mascot scores of each protein isoform were summarised at gene level (mean). The standard deviation as % of the mean per resulting gene was inspected, showing that variability for the vast majority of detected proteins was very small. Resulting tables were merged based on the gene symbol, mascot scores were log2 transformed and proteins were filtered based on being detected in all experimental samples in all experiments. Missing values were replaced by the minimum detected log2 value −1. Correlation between samples from the same experimental group was increased with this method and missing values greatly reduced. Data was imported into Perseus, analysed as described above, and visualised with ggplot2.

#### Data analysis ChIP-seq

ChIP-seq reads were aligned to the human reference genome build hg19 with bowtie (v1.1.1) (24). Peaks were called with MACS2 with default parameters and a matched input file (same ChIP protocol on the same cells) for transcription factors. In addition, the –broad argument was used for peak calling on histone marks (25). For heatmaps, the R package Genomation (version 1.30.0) was used (26). Peaks were sorted on either EVI1 enrichment or CTBP2 enrichment and widened symmetrically to 1000 bp on each side. These reference peaks were then used as windows to count signal in the BAM files of all tracks shown. The heatmaps are winsorised (“overexposed”) on (0,99), meaning all signal above the 99th percentile of peaks are the same colour. To quantify correlation in the heatmaps, DiffBind (27) (version 3.8.4) was used to count and normalise reads in all tracks within either EVI1 or CTBP2 peaks that were widened to 1000 bp on each side. Reads were normalized with default DiffBind parameters and correlation was calculated with linear regression on log10 transformed data and a pseudocount of 1. Correlation was visualised in ggplot2 with geom_hex. Individual loci were visualised with Gviz (version 1.42.1) (28). To annotate peaks to genes, DNAse hypersensitive sites significantly correlated with gene expression across diverse tissues were used (29) . If peaks did not overlap directly with a DNAse site in this dataset, they were annotated to the nearest protein-coding gene with ChIPPeakAnno (30) (version 3.32.0). To calculate fold changes for CTPB2 binding, Diffbind was used (same as above).

#### Data analysis RNA sequencing

RNAseq reads were quantified from fastq files directly using Salmon (v0.13.1) using hg38 RefGene database as an index. Transcript-level counts were aggregated to genes with tximport (v1.26.1) and differential gene expression analysis was performed with DEseq2 (v1.38.3). A significance cut-off of log2FC >1 and adjusted P-value <0.05 was used to determine the set of DE genes

#### AlphaFold

AlphaFold (v2.2.0, database updated to 08-01-2023) was run locally with 5x5 iterations and default parameters. Using Python, for each predicted relaxed multimer model, interaction residues were quantified with “interfaces”, “interfaces areaCutOff 0”, or “hbonds” in ChimeraX. The information for selecting residues was written to the log and extracted per structure. The resulting log files were important into R and combined into a single file with for each structure. The number of residues in each chain predicted to be interacting with each other was quantified and,if any, their identity was extracted. Data was visualised with ggplot2. 3D visualisations were done for the top ranked relaxed multimer models in ChimeraX.

### Xenotransplant models

#### Transplantation of SB1690CB cells into in NSG mice

Lentiviral supernatants of vectors expressing 4x PLDLS, 4x PLASS or empty controls, labelled with Emerald-GFP or mCherry were produced as described above. SB1690CB cells were transduced at multiplicity of infection (MOI) = 2 48 hours prior to injection into mice. The proportion of labelled cells was ascertained by flow cytometry prior to cells suspensions being combined to produce mixtures containing approximately 1:1 Emerald-GFP:mCherry labelled cells in three groups: Emerald-empty vector vs mCherry-empty vector, 4x PLASS Emerald vs 4x PLDLS mCherry, or 4x PLASS mCherry vs 4x PLDLS Emerald. For competitive engraftment, cell mixtures were administered into sublethally irradiated 8-12 week old female NSG mice (NOD/NSG/IL2Rgamma chain null) via intrabone injection of ∼1 x 106 filtered cells, suspended in 40 µL of sterile PBS (Gibco 10010056) containing 0.5% heat-inactivated FBS (Gibco 16140071). Mice were sacrificed after 10 weeks and suspensions of bone marrow cells prepared. After filtering, cells were washed twice and resuspended in PBS containing 2% FBS before staining in the presence of Fc block (Miltenyi 130-059-901) with anti-human CD45 Alexa700 (clone HI30, BioLegend cat #304024) and anti-mouse CD45.1 PE-Cy7 (clone A20, BioLegend cat #110730) antibodies and Hoechst 33258 (Thermo Fisher cat H3569). Cells were analysed using a BD FACSAria™ III Cell Sorter (BD Bioscience). The ratio of mCherry to Emerald-GFP expressing cells was determined as a proportion of human CD45 positive cells, following gating to exclude dead cells, doublets and murine CD45.1+ cells. Data was acquired using BD FACSDiva™ software v 8.0.1 (BD Bioscience) and tables of cell populations gated was exported from FACSDiva and prepared for analysis using the pandas data analysis library for Python. Frequency tables were visualised with ggplot. Xenograft experiments were performed with ethical approval under the Enver lab REC reference: 12/NW/0909. Care of animals, administration of cells, and animal sacrifice were performed in accordance with UK Home Office personal and project license requirements.

#### Transplantation of MUTZ3 into RAG2-/-γC-/- mice

Humanised scaffolds were prepared as described previously (31, 32). Briefly, mesenchymal stromal cells (MSCs) were isolated from healthy donor bone marrow, expanded in vitro and loaded onto biphasic calcium phosphate (BCP) particles. The MSC-BCPs were subcutaneously implanted in RAG2-/-γC-/- mice. MUTZ3 cells were stably transduced with a lentiviral luciferase vector (pLV-CMV Luc2 ires GFP) and subsequently with a retroviral 4xPLASS-Puro or 4xPLDLS-Puro vector as described above. Every 6-8 days, mice were anesthesized with isoflurance and I.P. injected with 100 μL 7.5 mM beetle luciferin (Promega). Mice were scanned in a third generation cooled GaAs intensified charge-coupled device camera, controlled by the Photo Vision software and analyzed with M3Vision software (all from Photon Imager; Biospace Laboratory) (33). Outgrowth data was normalized to day 0 and visualized with ggplot (geom_sina).

## Acknowledgements

The authors would like to thank Michael Vermeulen for flow cytometric analysis and cell sorting (FACS), the Proteomics facility at Erasmus MC for processing of mass spectrometry samples. We are endebted to Marijke Baltissen (Michiel Vermeulen group, Radboud University Nijmegen) for sharing protocols for nuclear protein extractions for mass spectrometry. For animal handling and care, we thank the animal facility at Erasmus MC. For the SB1690CB transplantation experiments, we would like to thank Preeta Datta (Tariq Enver lab, UCL Cancer Institute) who performed irradiation of mice, intrabone injections, care and sacrifice.

## Author contributions

Conceptualization, E.B., R.D., T.E., M.D.M.;

Investigation and methodology, D.P., M.H., D.B., W.N., J.G., L.S., J.A.A.D., R.W.J.G., E.B.;

Formal analysis and visualisation, D.P., R.M.L., R.H., J.A.A.D.;

Writing original draft, D.P., M.H., R.D.;

Writing review and editing, D.P., M.H, R. M. L., D.B., L.S., J.A.A.D., R.D.;

Supervision and project administration, R.D.

## Data and materials availability

ChIP-seq data is made available on GEO under accession number GSE236010. Raw mass spectrometry data and data for protein identification and quantification is submitted to the ProteomeXchange Consortium via the PRIDE partner repository with the data identifier PXD043333. All other data used for generating figures presented here (AlphaFold predicted structures, flow cytometry frequencies, luciferase output files and mass spec clustering and quantification) are available at Zenodo under accession 10.5281/zenodo.8354861. Custom scripts for AlphaFold quantifications and ChIP-seq correlation plots are available on Github (https://github.com/dorienpastoors/EVI1_CTBP2_PLDLSinhibitor). An overview of all data deposited associated with this manuscript per figure panel is available in Supplementary Table 1.

## References

1. E. Papaemmanuil et al., Genomic Classification and Prognosis in Acute Myeloid Leukemia. N Engl J Med 374, 2209–2221 (2016).

2. S. Ottema et al., Atypical 3q26/MECOM rearrangements genocopy inv(3)/t(3;3) in acute myeloid leukemia. Blood 136, 224–234 (2020).

3. S. Groschel et al., A single oncogenic enhancer rearrangement causes concomitant EVI1 and GATA2 deregulation in leukemia. Cell 157, 369–381 (2014).

4. L. Smeenk et al., Selective Requirement of MYB for Oncogenic Hyperactivation of a Translocated Enhancer in Leukemia. Cancer Discov 11, 2868–2883 (2021).

5. H. Yamazaki et al., A remote GATA2 hematopoietic enhancer drives leukemogenesis in inv(3)(q21;q26) by activating EVI1 expression. Cancer Cell 25, 415–427 (2014).

6. E. M. Bindels et al., EVI1 is critical for the pathogenesis of a subset of MLL-AF9-rearranged AMLs. Blood 119, 5838–5849 (2012).

7. R. A. Voit et al., A genetic disorder reveals a hematopoietic stem cell regulatory network co-opted in leukemia. Nat Immunol 24, 69–83 (2023).

8. V. Senyuk et al., The leukemia-associated transcription repressor AML1/MDS1/EVI1 requires CtBP to induce abnormal growth and differentiation of murine hematopoietic cells. Oncogene 21, 3232–3240 (2002).

9. I. Pinheiro et al., Prdm3 and Prdm16 are H3K9me1 methyltransferases required for mammalian heterochromatin integrity. Cell 150, 948–960 (2012).

10. K. Izutsu et al., The corepressor CtBP interacts with Evi-1 to repress transforming growth factor beta signaling. Blood 97, 2815–2822 (2001).

11. D. Ivanochko et al., Direct interaction between the PRDM3 and PRDM16 tumor suppressors and the NuRD chromatin remodeling complex. Nucleic Acids Res 47, 1225–1238 (2019).

12. E. A. Bard-Chapeau et al., EVI1 oncoprotein interacts with a large and complex network of proteins and integrates signals through protein phosphorylation. Proc Natl Acad Sci U S A 110, E2885–2894 (2013).

13. M. Kurokawa et al., The oncoprotein Evi-1 represses TGF-beta signalling by inhibiting Smad3. Nature 394, 92–96 (1998).

14. D. Spensberger, R. Delwel, A novel interaction between the proto-oncogene Evi1 and histone methyltransferases, SUV39H1 and G9a. FEBS Lett 582, 2761–2767 (2008).

15. D. Spensberger et al., Myeloid transforming protein Evi1 interacts with methyl-CpG binding domain protein 3 and inhibits in vitro histone deacetylation by Mbd3/Mi-2/NuRD. Biochemistry 47, 6418–6426 (2008).

16. S. Chakraborty, V. Senyuk, S. Sitailo, Y. Chi, G. Nucifora, Interaction of EVI1 with cAMP-responsive element-binding protein-binding protein (CBP) and p300/CBP-associated factor (P/CAF) results in reversible acetylation of EVI1 and in co-localization in nuclear speckles. J Biol Chem 276, 44936–44943 (2001).

17. H. Quentmeier, M. Zaborski, H. G. Drexler, The human bladder carcinoma cell line 5637 constitutively secretes functional cytokines. Leukemia Research 21, 343–350 (1997).

18. S. Meyer et al., A cross-linker-sensitive myeloid leukemia cell line from a 2-year-old boy with severe Fanconi anemia and biallelic FANCD1/BRCA2 mutations. Genes Chromosomes Cancer 42, 404–415 (2005).

19. C. C. Zhang, M. Kaba, S. Iizuka, H. Huynh, H. F. Lodish, Angiopoietin-like 5 and IGFBP2 stimulate ex vivo expansion of human cord blood hematopoietic stem cells as assayed by NOD/SCID transplantation. Blood 111, 3415–3423 (2008).

20. I. D. Karemaker, M. Vermeulen, ZBTB2 reads unmethylated CpG island promoters and regulates embryonic stem cell differentiation. EMBO Rep 19, (2018).

21. S. Tyanova, T. Temu, J. Cox, The MaxQuant computational platform for mass spectrometry-based shotgun proteomics. Nat Protoc 11, 2301–2319 (2016).

22. C. Stark et al., BioGRID: a general repository for interaction datasets. Nucleic Acids Res 34, D535–539 (2006).

23. G. Tsitsiridis, et al., CORUM: the comprehensive resource of mammalian protein complexes-2022. Nucleic Acids Res 51, D539–D545 (2023).

24. B. Langmead, C. Trapnell, M. Pop, S. L. Salzberg, Ultrafast and memory-efficient alignment of short DNA sequences to the human genome. Genome Biol 10, R25 (2009).

25. Y. Zhang et al., Model-based analysis of ChIP-Seq (MACS). Genome Biol 9, R137 (2008).

26. A. Akalin, V. Franke, K. Vlahovicek, C. E. Mason, D. Schubeler, Genomation: a toolkit to summarize, annotate and visualize genomic intervals. Bioinformatics 31, 1127–1129 (2015).

27. C. S. Ross-Innes et al., Differential oestrogen receptor binding is associated with clinical outcome in breast cancer. Nature 481, 389–393 (2012).

28. F. Hahne, R. Ivanek, Visualizing Genomic Data Using Gviz and Bioconductor. Methods Mol Biol 1418, 335–351 (2016).

29. N. C. Sheffield et al., Patterns of regulatory activity across diverse human cell types predict tissue identity, transcription factor binding, and long-range interactions. Genome Res 23, 777–788 (2013).

30. L. J. Zhu et al., ChIPpeakAnno: a Bioconductor package to annotate ChIP-seq and ChIP-chip data. BMC Bioinformatics 11, 237 (2010).

31. R. W. Groen et al., Reconstructing the human hematopoietic niche in immunodeficient mice: opportunities for studying primary multiple myeloma. Blood 120, e9–e16 (2012).

32. I. S. Nijhof et al., Preclinical Evidence for the Therapeutic Potential of CD38-Targeted Immuno-Chemotherapy in Multiple Myeloma Patients Refractory to Lenalidomide and Bortezomib. Clin Cancer Res 21, 2802–2810 (2015).

33. R. M. Reijmers et al., Targeting EXT1 reveals a crucial role for heparan sulfate in the growth of multiple myeloma. Blood 115, 601–604 (2010).

34. U. Vinatzer, J. Taplick, C. Seiser, C. Fonatsch, R. Wieser, The leukaemia-associated transcription factors EVI-1 and MDS1/EVI1 repress transcription and interact with histone deacetylase. Br J Haematol 114, 566–573 (2001).

35. C. E. Barnes, D. M. English, M. Broderick, M. O. Collins, S. M. Cowley, Proximity-dependent biotin identification (BioID) reveals a dynamic LSD1-CoREST interactome during embryonic stem cell differentiation. Mol Omics 18, 31–44 (2022).

36. P. J. Valk et al., The genes encoding the peripheral cannabinoid receptor and alpha-L-fucosidase are located near a newly identified common virus integration site, Evi11. J Virol 71, 6796–6804 (1997).

37. J. Jumper et al., Highly accurate protein structure prediction with AlphaFold. Nature 596, 583–589 (2021).

38. S. Palmer et al., Evi-1 transforming and repressor activities are mediated by CtBP co-repressor proteins. J Biol Chem 276, 25834–25840 (2001).

39. S. Lievens, F. Peelman, K. De Bosscher, I. Lemmens, J. Tavernier, MAPPIT: a protein interaction toolbox built on insights in cytokine receptor signaling. Cytokine Growth Factor Rev 22, 321–329 (2011).

40. P. Sontakke et al., Modeling BCR-ABL and MLL-AF9 leukemia in a human bone marrow-like scaffold-based xenograft model. Leukemia 30, 2064–2073 (2016).

41. R. Paredes et al., EVI1 oncoprotein expression and CtBP1-association oscillate through the cell cycle. Mol Biol Rep 47, 8293–8300 (2020).

42. J. Turner, M. Crossley, Cloning and characterization of mCtBP2, a co-repressor that associates with basic Kruppel-like factor and other mammalian transcriptional regulators. EMBO J 17, 5129–5140 (1998).

43. M. A. Blevins et al., CPP-E1A fusion peptides inhibit CtBP-mediated transcriptional repression. Mol Oncol 12, 1358–1373 (2018).

44. V. Kumar et al., Transcription corepressor CtBP is an NAD(+)-regulated dehydrogenase. Mol Cell 10, 857–869 (2002).

45. B. J. Hilbert, S. R. Grossman, C. A. Schiffer, W. E. Royer, Jr., Crystal structures of human CtBP in complex with substrate MTOB reveal active site features useful for inhibitor design. FEBS Lett 588, 1743–1748 (2014).

46. Y. Achouri, G. Noel, E. Van Schaftingen, 2-Keto-4-methylthiobutyrate, an intermediate in the methionine salvage pathway, is a good substrate for CtBP1. Biochem Biophys Res Commun 352, 903–906 (2007).

47. M. Sekiya et al., The transcriptional corepressor CtBP2 serves as a metabolite sensor orchestrating hepatic glucose and lipid homeostasis. Nat Commun 12, 6315 (2021).

48. S. S. Thio, J. V. Bonventre, S. I. Hsu, The CtBP2 co-repressor is regulated by NADH-dependent dimerization and possesses a novel N-terminal repression domain. Nucleic Acids Res 32, 1836–1847 (2004).

49. K. G. Quinlan et al., Role of the C-terminal binding protein PXDLS motif binding cleft in protein interactions and transcriptional repression. Mol Cell Biol 26, 8202–8213 (2006).

