## Supplemental Figures and Table Legend for "Oncogene EVI1 Drives Acute Myeloid Leukemia Via a Targetable Interaction with CTBP2"

Dorien Pastoors *et al.*

**This PDF file includes:**

Figs. S1 to S5

**Other Supplementary Materials for this manuscript include the following:**

Data S1. Supplementary table – Description of sheets in supplementary file

**Figure S1: Supplement to figure 1 (Pt1)**

Fig S1A. EVI1 IP followed by CTBP2 Western blot (left panel) or vice versa (right panel) in MUTZ3 whole-cell lysates.

Fig S1B. Full version of Fig 1B, including all the protein labels and groups of proteins removed because they are likely contaminants.

Fig S1C. Outline of EVI1 pulldown experiments with EVI1-Biotag and BirA overexpression in NFS78

Fig S1D. MS enrichment of proteins in streptavidin IP of BirA+ EVI1-Biotag versus BirA alone in murine cell line NFS78. (Significance cut-off: log2FC > 1 and p-value < 0.05 (n = 4 for each group))

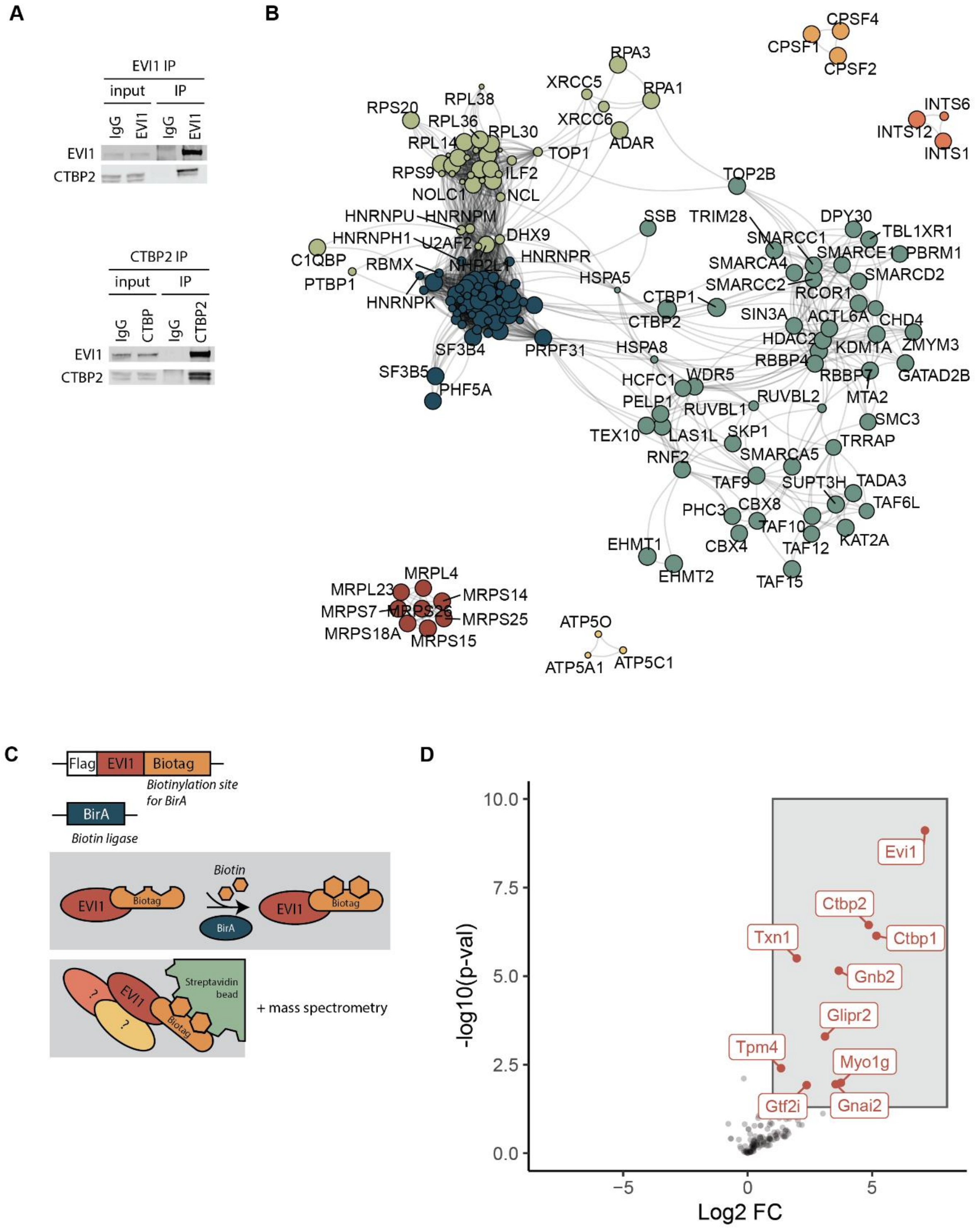
A
EVI1 IP
input
IP
IgG
EVI1
IgG
EVI1
EVI1
CTBP2
CTBP2 IP
input
IP
IgG
CTBP
IgG
CTBP2
EVI1
CTBP2
B
CPSF4
CPSF1
CPSF2
INTS6
INTS12
INTS1
RPA3
RPA1
XRCC5
XRCC6
ADAR
RPL38
RPS20
RPL36
RPL30
RPL14
TOP1
RPS9
ILF2
NOLC1
NCL
HNRNPU
HNRNPM
HNRNPH1
U2AF2
DHX9
C1QBP
RBMX
NHP2L1
HNRNPR
PTBP1
HNRNPK
SF3B4
PRPF31
SF3B5
PHF5A
TOP2B
SSB
TRIM28
DPY30
TBL1XR1
SMARCC1
SMARCE1
PBRM1
SMARCA4
SMARCD2
SMARCC2
RCOR1
HSPA5
CTBP1
CTBP2
SIN3A
ACTL6A
CHD4
HDAC2
ZMYM3
KDM1A
HSPA8
RBBP4
GATAD2B
RBBP7
HCFC1
WDR5
MTA2
PELP1
RUVBL1
RUVBL2
SMC3
TEX10
LAS1L
SKP1
TRRAP
RNF2
SMARCA5
TAF9
SUPT3H
TADA3
PHC3
CBX8
TAF6L
TAF10
CBX4
KAT2A
TAF12
EHMT1
EHMT2
TAF15
MRPL4
MRPL23
MRPS14
MRPS7
MRPS26
MRPS25
MRPS18A
MRPS15
ATP5O
ATP5A1
ATP5C1
C
Flag
EVI1
Biotag
Biotinylation site for BirA
BirA
Biotin ligase
Biotin
EVI1
Biotag
BirA
EVI1
Biotag
?
EVI1
?
Biotag
Streptavidin bead
+ mass spectrometry
D
10.0
7.5
5.0
2.5
0.0
-log10(p-val)
Evi1
Ctbp2
Txn1
Ctbp1
Gnb2
Glipr2
Tpm4
Myo1g
Gtf2i
Gnai2
−5
0
5
Log2 FC

S1

**Figure S2. Supplement to Fig.1 (Pt2)**

Fig. S2A. Quantification of the heatmap in panel Fig. S2B. For each track, the normalised CTBP2 signal is plotted versus the normalised reads in the indicated ChIP-seq-tracks, for all CTBP2 peaks with a window of ±1000bp. Correlation coefficients and linear regression equations are shown for log10-transformed data with a pseudo count of 1.

Fig. S2B. Heatmap ranked on ChIP-seq CTBP2 signal (leftmost panel) showing signal intensity of indicated ChIP-seq tracks within CTBP2 peaks

Fig. S2C. Western blot of inducible EVI1 knock-down in a clone derived from MOLM1 performed in the same experiment as the ChIP-seq in Fig. 1E (48 hrs of doxycycline)

A

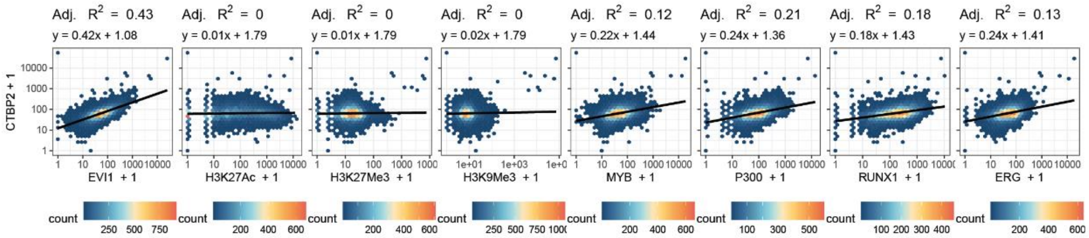
Adj. R2 = 0.43
y = 0.42x + 1.08
Adj. R2 = 0
y = 0.01x + 1.79
Adj. R2 = 0
y = 0.01x + 1.79
Adj. R2 = 0
y = 0.02x + 1.79
Adj. R2 = 0.12
y = 0.22x + 1.44
Adj. R2 = 0.21
y = 0.24x + 1.36
Adj. R2 = 0.18
y = 0.18x + 1.43
Adj. R2 = 0.13
y = 0.24x + 1.41
CTBP2 + 1
EVI1 + 1
H3K27Ac + 1
H3K27Me3 + 1
H3K9Me3 + 1
MYB + 1
P300 + 1
RUNX1 + 1
ERG + 1
count

B

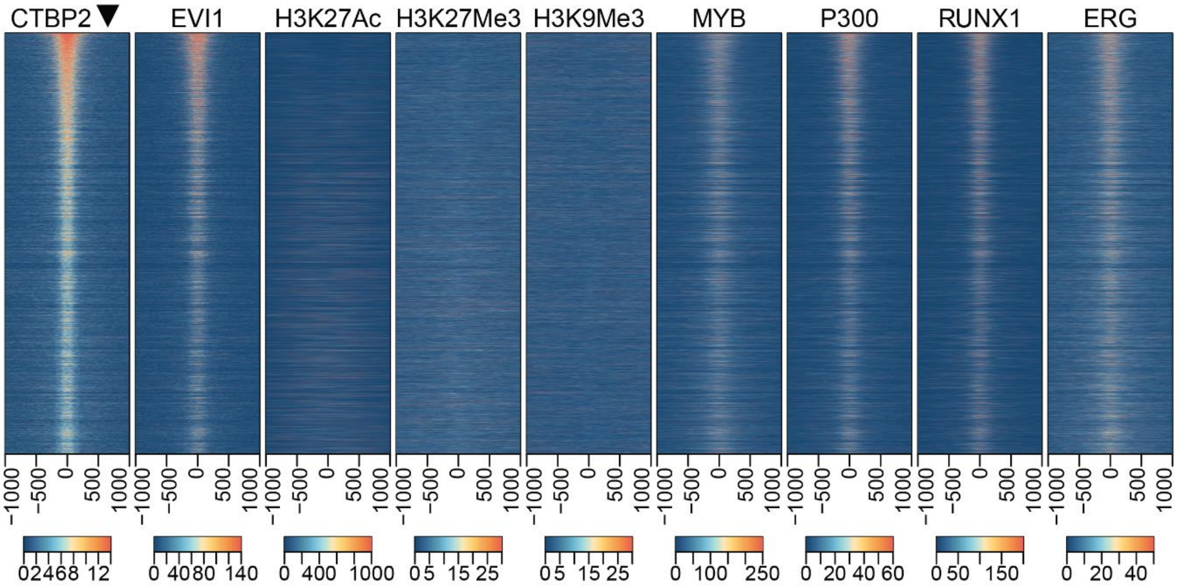
CTBP2
EVI1
H3K27Ac
H3K27Me3
H3K9Me3
MYB
P300
RUNX1
ERG

C

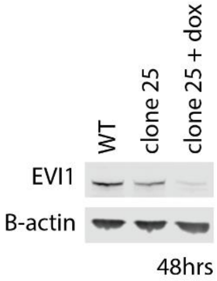
WT
clone 25
clone 25 + dox
EVI1
B-actin
48hrs

**Figure S3: Supplement to figure 2-3**

Fig. S3A. Schematic outline of the MAPPIT assay to measure protein-protein interaction (here for EVI1 and CTBP2). The MAPPIT receptor is based on the EPO receptor, which harbors an intracellular domain that can be phosphorylated by JAK in normal JAK-STAT signalling (1). In the modified MAPPIT receptor, the intracellular domain fused to EVI1 and the Tyr phosphorylation sites are mutated. Gp130 is fused to CTBP2 (2). Because EVI1-CTBP2 can interact with each other, CTBP2 is recruited to the EPO-EVI1 receptor. In this way, when signal is transmitted by ligand binding (2), JAK will phosphorylate tyrosines on CTBP2-gp130 to subsequently activate STAT3 (3). Phosphorylated STAT3 is transported into the nucleus where it will activate a STAT3 responsive luciferase reporter (4).

Fig. S3B. MAPPIT assay on HEK293T cells of EVI1 and CTBP2, with amino acid (AA) segments of varying sizes around the PLDLS site (100AA is ±100AA) (n=1 in triplicate).

Fig. S3C. MAPPIT assay on HEK293T cells (n=6, in triplicate) of ZEB2 and CTBP2, with inhibitor of EVI1 and CTBP interaction. Reporter induction normalised to –EPO (not induced) and maximum induction with RNF positive control. Significance with two-way ANOVA (comparison PLDLS vs PLASS + EPO shown).

Fig. S3D. IP EVI1-WB CTBP1/2 on MUTZ3 cells, stained for EVI1 and CTBP2 with 4x PLASS-FLAG or 4x PLDLS-FLAG dox-inducible overexpression construct.

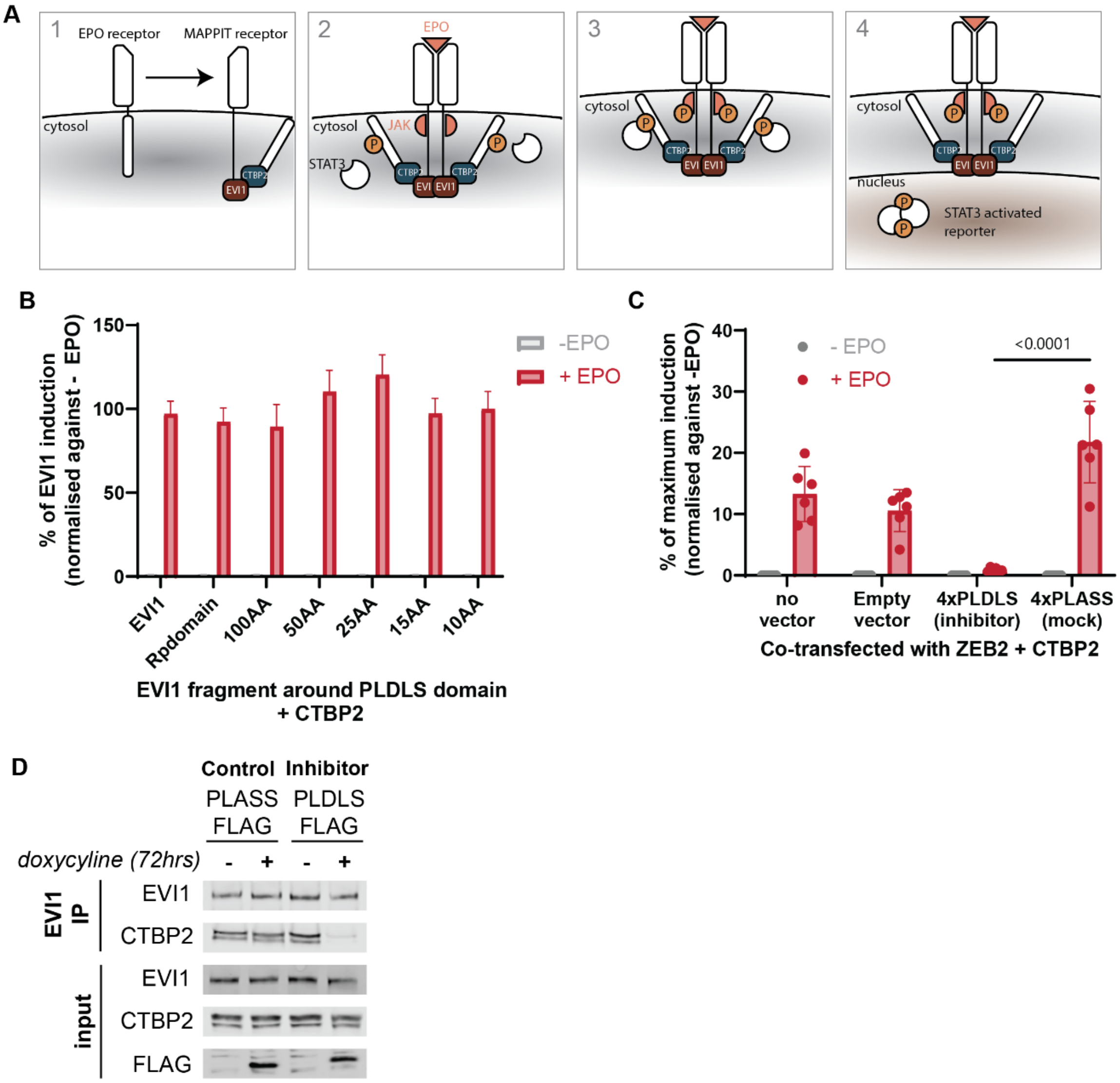
A
1 EPO receptor MAPPIT receptor
cytosol
EVI1
CTBP2
2 EPO
cytosol JAK
STAT3
3
cytosol
4
cytosol
nucleus
STAT3 activated reporter
B
% of EVI1 induction (normalised against - EPO)
-EPO
+ EPO
EVI1
Rpdomain
100AA
50AA
25AA
15AA
10AA
EVI1 fragment around PLDLS domain + CTBP2
C
% of maximum induction (normalised against -EPO)
- EPO
+ EPO
<0.0001
no vector
Empty vector
4xPLDLS (inhibitor)
4xPLASS (mock)
Co-transfected with ZEB2 + CTBP2
D
Control
Inhibitor
PLASS FLAG
PLDLS FLAG
doxycyline (72hrs)
EVI1 IP
input
EVI1
CTBP2
FLAG

**Figure S4: Supplement to Figure 3**

Fig. S4A. Volcano plot of differential CTBP2 binding site occupancy (by ChIP-seq) analysis (DiffBind) in 4x PLASS vs 4x PLDLS -treated MUTZ3 cells (5281 peaks with FDR < 0.05; n = 2).

Fig. S4B. Fold-change of CTBP2 peaks in response to 4x PLDLS treatment in MUTZ3. Peaks are grouped according to whether they overlap with EVI1 peaks (n = 4026), or not (n =1608).

Fig. S4C. Significantly differentially expressed (DE) genes (adj. P-value <0.05 and absolute fold-change > 2) upon 4x PLDLS treatment in MUTZ3 cells (n = 2, 72 hrs post-TD, 162 genes in total). The fraction of DE genes that are bound by CTBP2 are highlighted in dark. Peak-to-gene annotation is based firstly on whether the peak directly overlaps with a putative regulatory element associated to gene expression, and if it does not overlap with such an element, to the nearest protein-coding gene.

Fig. S4D. CTBP2 peaks are ranked (x-axis) based on their enrichment over input (y-axis). The top-30 peaks annotated to genes from figure C based on highest CTBP2 binding are labelled. A. The distribution of all PLDLS-target-gene-annotated-peaks is visualised above the axis. 342 peaks could be annotated to a upregulated PLDLS target gene (88 unique genes in total, on average 3.9 peaks per gene) and 47 to a downregulated PLDLS target gene (18 unique genes in total, or 2.6 peaks per gene on average).

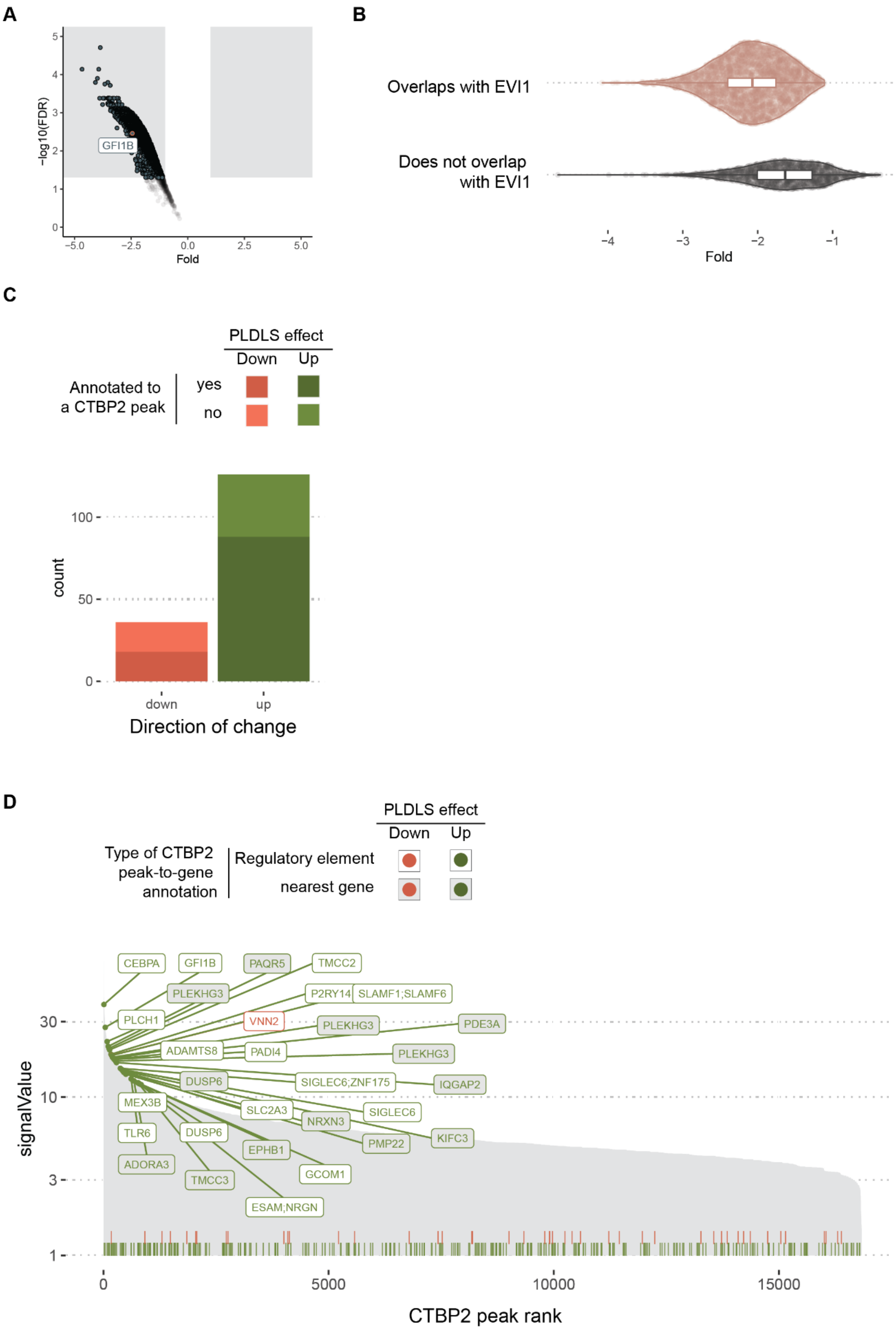
A
−log10(FDR)
GFI1B
Fold
−5.0
−2.5
0.0
2.5
5.0
B
Overlaps with EVI1
Does not overlap with EVI1
Fold
−4
−3
−2
−1
C
PLDLS effect
Down
Up
Annotated to a CTBP2 peak
yes
no
count
0
50
100
down
up
Direction of change
D
PLDLS effect
Down
Up
Type of CTBP2 peak-to-gene annotation
Regulatory element
nearest gene
CEBPA
GFI1B
PAQR5
TMCC2
PLEKHG3
P2RY14
SLAMF1;SLAMF6
PLCH1
VNN2
PLEKHG3
PDE3A
ADAMTS8
PADI4
PLEKHG3
DUSP6
SIGLEC6;ZNF175
IQGAP2
MEX3B
SLC2A3
SIGLEC6
NRXN3
TLR6
DUSP6
KIFC3
PMP22
EPHB1
ADORA3
TMCC3
GCOM1
ESAM;NRGN
signalValue
1
3
10
30
0
5000
10000
15000
CTBP2 peak rank

**Figure S5. Supplement to Figure 5**

Fig S5A. Flow-cytometric analysis of input and 5 mice transplanted with SB1690 with Empty vector-Emerald and Empty vector -mCherry mixed in a 1:1 ratio.

Fig S5B. Flow-cytometric analysis of input and 7 mice transplanted with SB1690 with PLASS-Emerald and PLDLS -mCherry mixed in a 1:1 ratio.

Fig S5C. Luminescence scans of mice transplanted with MUTZ3-Luciferase +4xPLASS or MUTZ3-Luciferase +4x PLDLS vectors at indicated days after transplantation

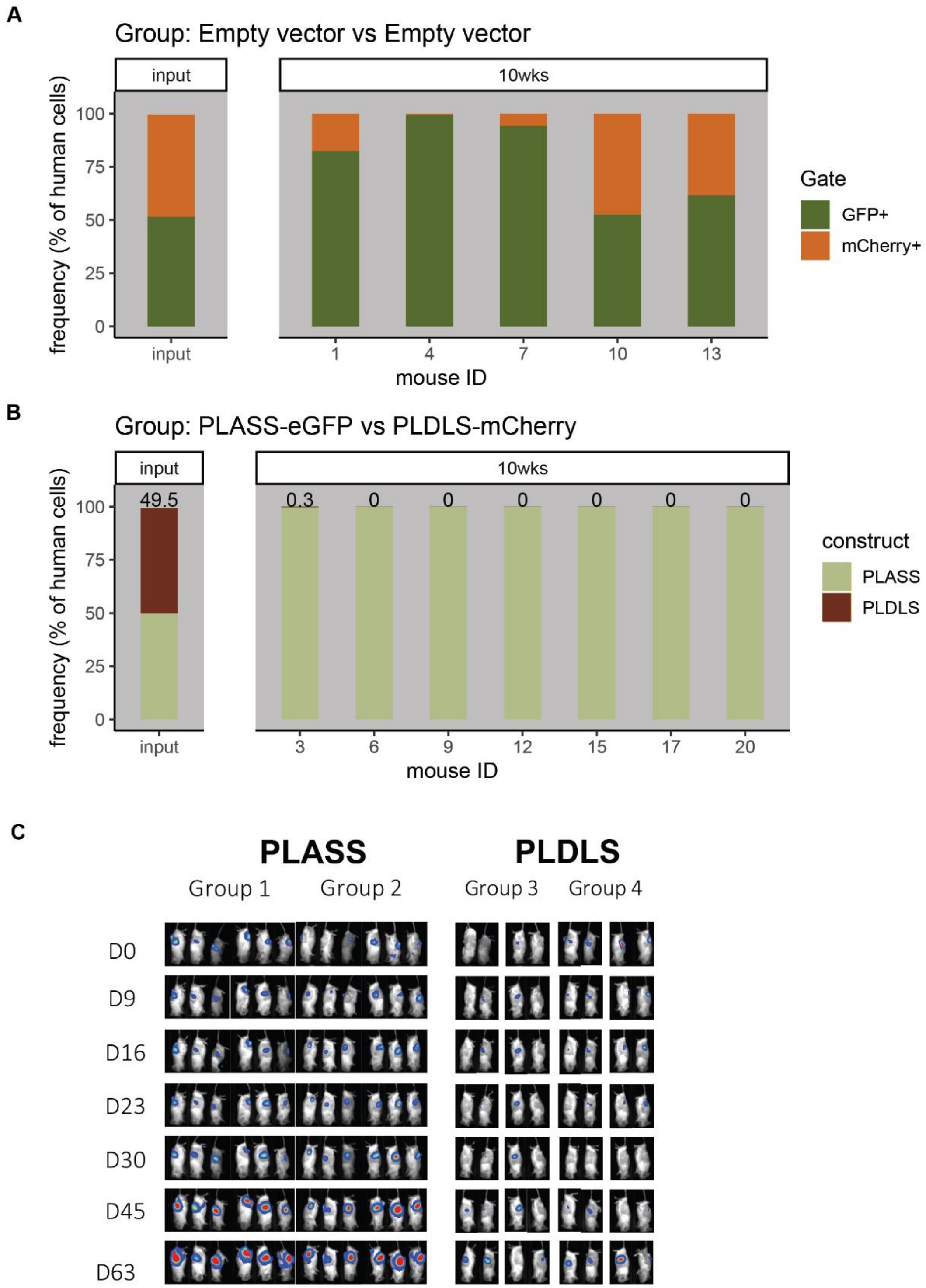
A
Group: Empty vector vs Empty vector
input
10wks
frequency (% of human cells)
100
75
50
25
0
input
1
4
7
10
13
mouse ID
Gate
GFP+
mCherry+
B
Group: PLASS-eGFP vs PLDLS-mCherry
input
10wks
49.5
0.3
0
0
0
0
0
0
frequency (% of human cells)
100
75
50
25
0
input
3
6
9
12
15
17
20
mouse ID
construct
PLASS
PLDLS
C
PLASS
PLDLS
Group 1
Group 2
Group 3
Group 4
D0
D9
D16
D23
D30
D45
D63

**Data S1. (SupplementaryTable.xlsx)**

Supplementary Table 1 contains the following sheets:

- Oligos > all used oligos in the study
- GEO_perfigure> description of all sequencing data used per figure panel (all available on GEO; mostly ChIP-seq and some RNAseq)
- Chipseq_antibodies_protocols > all antibodies per chip track and the used protocol per track
- DataAvailability_PerFigure > description of all datasets associated with this publication deposited in public repositories (GEO, ProteomeXchange, Github and ZENODO).
- ColonyAssays_WithinGroup >
    - Results of two-way ANOVA on within-group comparisons for Fig 4B.
    - Results of two-way ANOVA on within-group comparisons for Fig 5A.
